# Spatial task instructions and global activation trends influence functional modularity in the cortical reach network

**DOI:** 10.1101/2025.03.20.644329

**Authors:** L. Musa, A. Ghaderi, Y. Chen, J.D Crawford

## Abstract

Humans can be instructed to ignore visual cues or use them as landmarks for aiming movements (Musa et al. 2024), but it is not known how such allocentric cues interact with egocentric target codes and general planning activity to influence cortical network properties. To answer these questions, we applied graph theory analysis (GTA) to a previously described fMRI dataset (Chen et al. 2014). Participants were instructed to reach toward targets defined in egocentric or landmark-centered (allocentric) coordinates. During *Egocentric* pointing, cortical nodes clustered into four bilateral modules with correlated BOLD signals: a superior occipital-parietal / somatomotor module, an inferior parietal / lateral frontal module, a superior temporal / inferior frontal module, and an inferior occipital-temporal / prefrontal module. The *Allocentric* task showed only three modules, in part because inferior occipital nodes were incorporated into the superior occipital-parietal / somatomotor module. Both tasks engaged local (within module) and global (between module) cortical hubs, but the *Allocentric* task recruited additional hubs associated with allocentric visual codes and ego-allocentric integration. Removing reach-related activation trends reduced global synchrony and increased clustering, specifically diminishing dorsoventral coupling in the allocentric task. Cross-validated decoding confirmed that modularity provided the best predicter of task type and suggest that temporal / parietal modules spanning prefrontal cortex play an important role in task instruction. These results demonstrate that activation trends related to motor plans influence global network integration, whereas task instructions influence intermediate / local network properties, such as the modular integration and hub recruitment observed in our *Allocentric* task.

**Highlights:**

- The study explores how egocentric and allocentric cues affect cortical networks.
- Graph theory analysis (GTA) was applied to fMRI data from pointing tasks.
- Egocentric pointing formed four cortical modules; allocentric formed three.
- Allocentric tasks recruited additional hubs for dorsal-ventral integration.
- Removing reach-related activation trends reduced global synchrony in the allocentric task.

## 1. Introduction

The neural mechanisms that process spatial information are central to our ability to interact with the environment. Typically, this involves an interaction between specific task parameters, sensory inputs, and motor plans. For example, humans can be instructed to reach toward visual objects using egocentric cues (i.e., location relative to the eyes, head, or body) or allocentric cues, i.e., location relative to visual landmarks (Byrne & Crawford, 2010a, 2010b; Lemay et al., 2004; Musa et al., 2024). It is thought that the visual system processes egocentric versus allocentric information through different cortical ‘streams’ (Milner & Goodale, 2006), but it is not known how task specific task instructions interacts with these signals and motor planning signals to influence segregation and integration of this information at whole-brain, intermediate, and local levels. The present study investigated this question by identifying whole brain network parameters, modules, hubs, and mechanisms for cross-module integration, contrasting networks derived from neuroimaging data collected during egocentric versus landmark-centered pointing tasks.

Neuropsychological studies suggest that the dorsal visual stream, running from occipital to parietal cortex, is associated with egocentric coding of vision-for-action, whereas the ventral stream is associated with allocentric coding and perception (Goodale & Milner 1912; Milner & Goodale, 2006; Schenk & McIntosh, 2010). This distinction was supported by subsequent neuroimaging studies (e.g. Committeri et al. 2004; Galati, 2000; Zaehle et al., 2007), although the ventral stream is engaged when a memory delay precedes action (Budisavljevic, 2018; Goodale et al., 2005; Vesia & Crawford, 2012; Khan et al., 2013).

In a more recent neuroimaging study, Chen et al. (2014) presented participants with both reach targets and visual landmarks and either instructed them to ignore the landmark (*Egocentric* task) or remember target location relative to the landmark (*Allocentric* task). They then shifted the landmark during a memory interval, to dissociate egocentric and allocentric coordinates. Their imaging results suggest that the task instruction reproduced a dorsoventral dissociation for egocentric / allocentric coding during the memory interval. Further, directional selectivity for target location was evident in both egocentric and allocentric frames, with areas like the inferior occipital gyrus (IOG) and superior occipital gyrus (SOG) showing preferential activity for egocentric direction coding, while the inferior temporal gyrus (ITG) was more selective for allocentric coding, requiring an interaction between target and landmark information in the ventral stream.

Other studies have emphasized the need to integrate information across these streams for real-world behavior (Prime et al., 2007; Freud et al., 2018, Baltaretu, et al., 2020). For example, various behavioral studies have shown that allocentric cues influence reach, even when participants are instructed to ignore visual landmarks (Byrne & Crawford, 2010; Fiehler et al., 2014). Based on the two visual streams hypothesis, this would ultimately require the incorporation of ventral stream allocentric target information into the dorsal stream processes associated with action planning (Chen et al., 2014). For example, precuneus, supplementary motor, and dorsal premotor cortex were selectively activated when participants are cued to reach toward a target location that was stored and remembered relative to a landmark (Chen et al., 2018). Consistent with this, neurophysiological studies have shown that prefrontal visual, memory, and motor signals integrates egocentric and allocentric signals for action (Bharmauria et al. 2020, 2021; Schutz et al., 2023). Overall, these results support the classic dorsal/ventral distinction for ego/allocentric coding during memory-guided reach but suggest that multiple cortical areas may be involved in integrating this information for action.

Finally, when participants are instructed to perform landmark-centred reach, the parietofrontal networks for egocentric action planning and execution are recruited, along with a progressive rise in BOLD activation across large swaths of cortex (Chen et al. 2014, 2018). The latter phenomenon has been observed in various sensorimotor imaging studies (Medendorp et al., 2003, Cappadocia et al. 2017) and might be attributed to recurrent feedback from the motor system (Gallivan & Culham, 2015; Blohm et al., 2019; Monaco et al. 2024). However, the possible functional role of this widespread, progressive BOLD activation for signal integration has not been considered, to our knowledge.

These findings suggest that understanding the interaction between egocentric and allocentric codes requires an integrated, whole-brain approach, based not just on regions of interest, but also on how information is propagated across these areas. Recent advances in network science, for example through the application of graph theory analysis (GTA), provide a valuable framework for modeling the complex neural interactions of signals within and between different brain regions (Sporns, 2011). Specifically, GTA provides a means of formalizing large-scale patterns of ‘functional connectivity’ (i.e., correlations between time series data derived from local ‘nodes’) at the whole-brain level. This technique is employed most often in conjunction with resting-state neuroimaging data (Braun et al., 2012) but can also be used to derive task-related networks from event-related data (Ghaderi et al., 2023; Tomou et al., 2024). This provides an objective means to quantify network parameters at the whole brain level (i.e., global parameters), intermediate levels (i.e., modular clusters of nodes with similar signals), and local levels (i.e.,Hubs: nodes that correlate strongly with other important nodes).

The current study investigated the influence of an allocentric (vs. egocentric) instruction on the cortical reach network by applying graph theory analysis to the dataset collected by Chen et al. (2014). Specifically, we aimed to contrast the cortical networks for egocentric vs. allocentric memory-guided reach, based on global network parameters for segregation and integration of information, objectively identified modules, and important hubs for sharing local and global task information. Based on previous findings by Chen et al. (2014, 2018) and others, we hypothesized that 1) removing overall whole-brain trends related to sensorimotor planning and execution would affect global and intermediate network properties, whereas 2) task instruction would influence intermediate and local levels, i.e., increased communication between dorsal and ventral visual network modules, and recruitment of network hubs associated with specific task requirements. Finally, we evaluated the capacity of these networks to predict behaviour by decoding reach task (egocentric vs. allocentric) from global and intermediate network parameters.

## 2. Methods

### 2.1 Data Source

This study involved a secondary analysis of data that were previously published in a univariate ‘region of interest’ study (Chen et al. 2014). The dataset consisted of neuroimaging data of twelve right-handed participants (8 females, 4 males; aged 23–40 years). All participants had normal or corrected-to-normal vision and were free of any known neuromuscular impairments.

### 2.2 Experimental Stimuli and Apparatus

Complete details of the experiment are provided in Chen et al. 2014; here we repeat the main points necessary to understand the task and analysis. The experiment was conducted at York University’s neuroimaging center using a 3-T Siemens Magnetom TIM Trio MRI system. A 12-channel head coil (6 channels posterior, 4-channel flex coil anterior) was used for data acquisition. Experiments were performed in near-complete darkness (participants reported that they could not see the workspace or hand) to minimize extraneous visual cues. Visual stimuli consisted of 3 mm LED lights controlled by fiber optic cables embedded in a custom board mounted above the participant’s abdomen. The platform was adjustable for optimal comfort, visibility and reach accessibility, and affixed to the MRI bed. A touch screen (Keytec) recorded reaching endpoints, while gaze was tracked using an iView X eye-tracking system with an MRI-compatible Avotec Silent Vision system. The head was slightly tilted (20°) to allow direct viewing of the stimuli. Participants wore headphones to receive audio instructions for the upcoming trial and before pointing.

### 2.3 Stimulus Specifications

Visual stimuli were color-coded: yellow for gaze fixation, green/red for reach targets, and blue for landmarks (Figure 1). Reach targets and landmarks were located to the left and right of central gaze fixation points (Figure 1). Target and landmark positions varied horizontally (with a range of 1-2° visual angle from target), with 40 possible configurations. A ‘mask’ consisting of 20 LEDs was positioned above and below the visual stimuli, vertically aligned between the target and landmark. The mask was presented to minimize aftereffects from the target and landmark stimuli, ensuring memory encoding rather than afterimage reliance. Light intensity was set to a level that was clearly visible for the participant but did not illuminate the hand or workspace.

**Figure 1:**
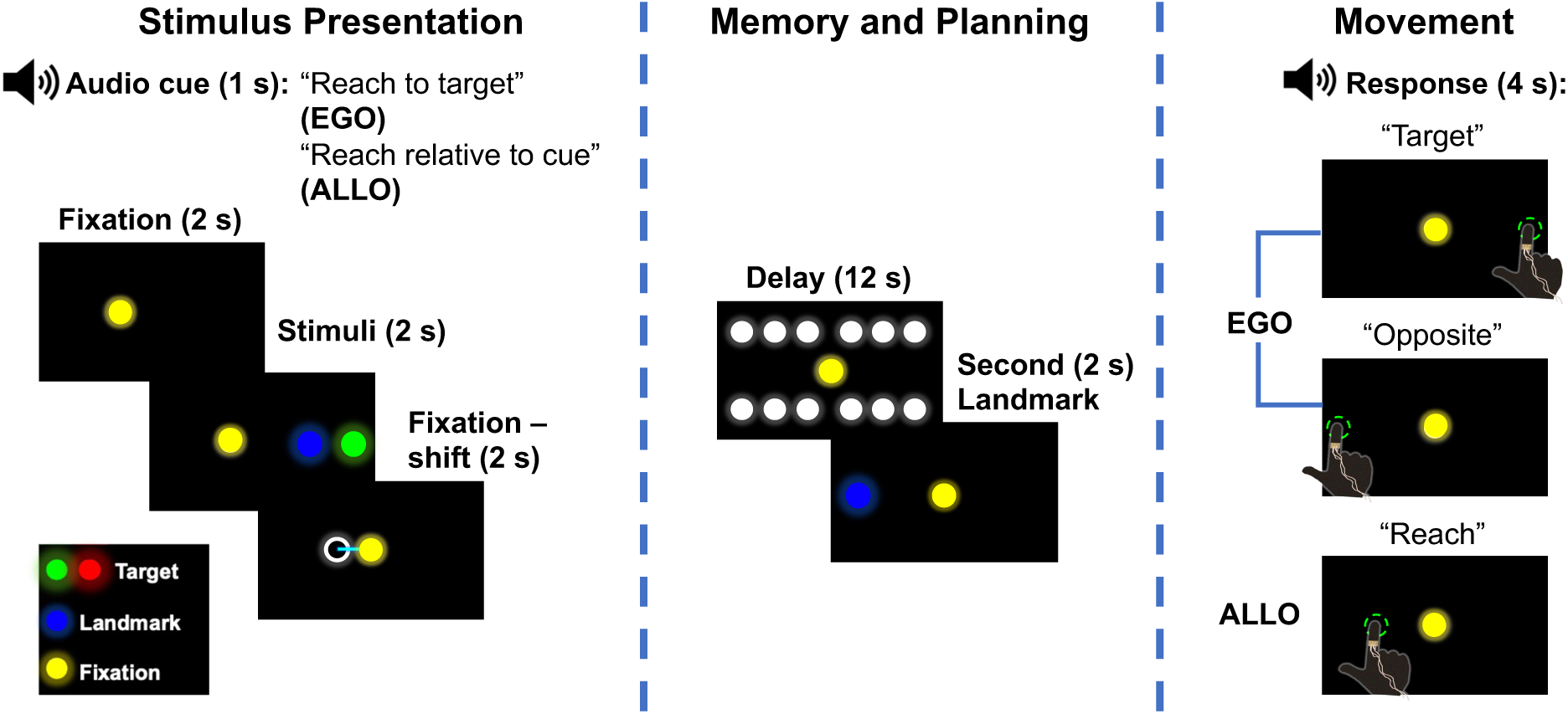
Task. Experimental paradigm, divided into three sequential phases: stimulus presentation (6 seconds total), memory and planning (14 seconds total), and movement (4 seconds). Audio task instructions preceded stimulus presentation and differed based on the type of task, Egocentric – participants were instructed to remember the target location only, Allocentric – participants were instructed to remember the target location relative to the landmark. Black boxes represent the stimulus pad / touch screen and the dots represent the fixation point (yellow), target (red or green), landmark (blue) and mask (white). With the exception of the initial task audio instructions, the stimuli presented during the stimulus presentation phase and the memory and movement planning phases are the same for the Egocentric and Allocentric task. However, during the movement planning phase participants also heard differing instructions to execute different types of movements based on the final task relative audio instructions, which prompted participants to perform a pro or anti point in the Egocentric task or point to the remembered position relative to the second landmark. See text for details.

### 2.4 Experimental Paradigm and Timing

The experiment featured two primary tasks: *Egocentric* and *Allocentric* (Figure 1). A third non-spatial *control* task (not shown) was used in the original univariate analysis (Chen et al. 2014) but was not required here. Each trial began with an audio cue indicate the task: *Egocentric*, where participants were instructed to remember where they saw the object, *Allocentric* task, where participants were instructed to remember the location of the object relative to the landmark. Participants then looked at the gaze fixation point, followed by a 2-second display of the target and landmark. The fixation stimulus then shifted slightly left or right so that it could not be used as an allocentric landmark. This was followed by a 12-second memory delay, during which the mask was presented. Afterward, the landmark reappeared either at the same or different location, and participants were cued to perform the reach toward the target (in an egocentric or allocentric coordinates, as initially instructed). Since the landmark appeared in the opposite hemifield 50% of the time, tasks were equalized by providing a *Pro/Anti* instruction before *Egocentric* reaches, i.e., to reach toward or opposite (mirrored across midline) to remembered target location. A 16-second inter-trial interval was provided to allow for hemodynamic baseline recovery. Participants practiced the tasks until they did them correctly, one day before recordings.

The design incorporated three factors: task (*Egocentric, Allocentric*), target position relative to gaze (left or right), and target position relative to the landmark (left or right). This resulted in 12 counterbalanced conditions across six experimental runs, each consisting of 18 trials. Note that the distribution of stimuli (targets and landmarks) was essentially the same for each task, only the instruction differed.

### 2.5 Imaging Parameters

Functional data were acquired with an echo-planar imaging (EPI) sequence (TR = 2000 ms, TE = 30 ms, FA = 90°, FOV = 192 x 192 mm, matrix = 64 x 64, slice thickness = 3.5 mm, 35 slices, angled at 25°, ascending interleaved). T1-weighted anatomical images were acquired using an MPRAGE sequence (TR = 1900 ms, TE = 2.52 ms, inversion time = 900 ms, FA = 9°, FOV = 256 x 256 mm, voxel size = 1 x 1 x 1 mm³).

### 2.6 Preprocessing

Data preprocessing was performed using FSL (FMRIB Software Library). The first two volumes of each run were discarded to account for saturation effects. Slice time correction, motion correction, and high-pass temporal filtering (to remove low-frequency noise) were applied. Motion correction was performed using FSL’s MCFLIRT. Functional data were co-registered to the T1 anatomical scan using FLIRT (FMRIB’s Linear Image Registration Tool), and then normalized to MNI space. Spatial smoothing was applied using a Gaussian kernel with a 6-mm full-width half-maximum (FWHM).

### 2.7 Data Analysis

For individual analysis, a General Linear Model (GLM) was constructed using FSL’s FEAT (FMRIB’s Expert Analysis Tool) for each participant, including task-specific predictors for different phases of the experiment (eye movement, target presentation, delay, landmark, and response phases). Six motion correction parameters were included as regressors of no interest to control for head movement. Task conditions included EGOCENTRIC and ALLOCENTRIC reaches and the **color** task, each with target positions defined relative to gaze or landmark.

#### 2.7.1 Voxelwise Analysis

Voxelwise analysis was conducted using a group-level mixed-effects analysis with FLAME (FMRIB’s Local Analysis of Mixed Effects). Group contrasts were performed using higher-level FEAT analysis in FSL. The following contrasts were tested:

1. **EGOCENTRIC vs. Baseline (Delay Phase)**: Areas coding target location during the delay phase in the **EGOCENTRIC** task.
2. **ALLOCENTRIC vs. Baseline (Delay Phase)**: Areas coding target location during the delay phase in the **ALLOCENTRIC** task.
3. **EGOCENTRIC Left vs. EGOCENTRIC Right (Delay Phase)**: Directional selectivity within the **EGOCENTRIC** task (left vs. right relative to gaze).
4. **ALLOCENTRIC Left vs. ALLOCENTRIC Right (Delay Phase)**: Directional selectivity within the **ALLOCENTRIC** task (left vs. right relative to landmark).
5. **Reach Left vs. Reach Right (Response Phase)**: Directional selectivity in the **EGOCENTRIC** and **ALLOCENTRIC** reach tasks based on the actual movement direction.

#### 2.7.2 Statistical Corrections

To correct for multiple comparisons, we applied cluster-based correction using FSL’s randomise tool (1000 permutations) to estimate the significance of contiguous clusters of activation. The minimum cluster size for inclusion was 11 voxels (297 mm³). A Bonferroni correction was applied to post-hoc paired-sample t-tests to control for Type I error across multiple comparisons, with a corrected p-value threshold of 0.0167.

### 2.8 Regions of Interest and Time Series Analysis

In this study, we chose 200 regions of interest (ROIs) as nodes, each defined as a 6-mm diameter spherical region based on the centroid coordinates of the Schaefer-Yeo atlas (Schaefer et al., 2018) (Figure 2A). Average timeseries for entire runs were then extracted at each ROI (Figure 2B) using the fslmeants function. Each run (6 in total) consisted of 6 egocentric task trials and 6 allocentric task trials, which were analysed independently by isolating the time series from the start of the stimulus presentation phase until the end of the delay epoch (18 s in total), creating timeseries as shown in the example in Figure 2C.

**Figure 2:**
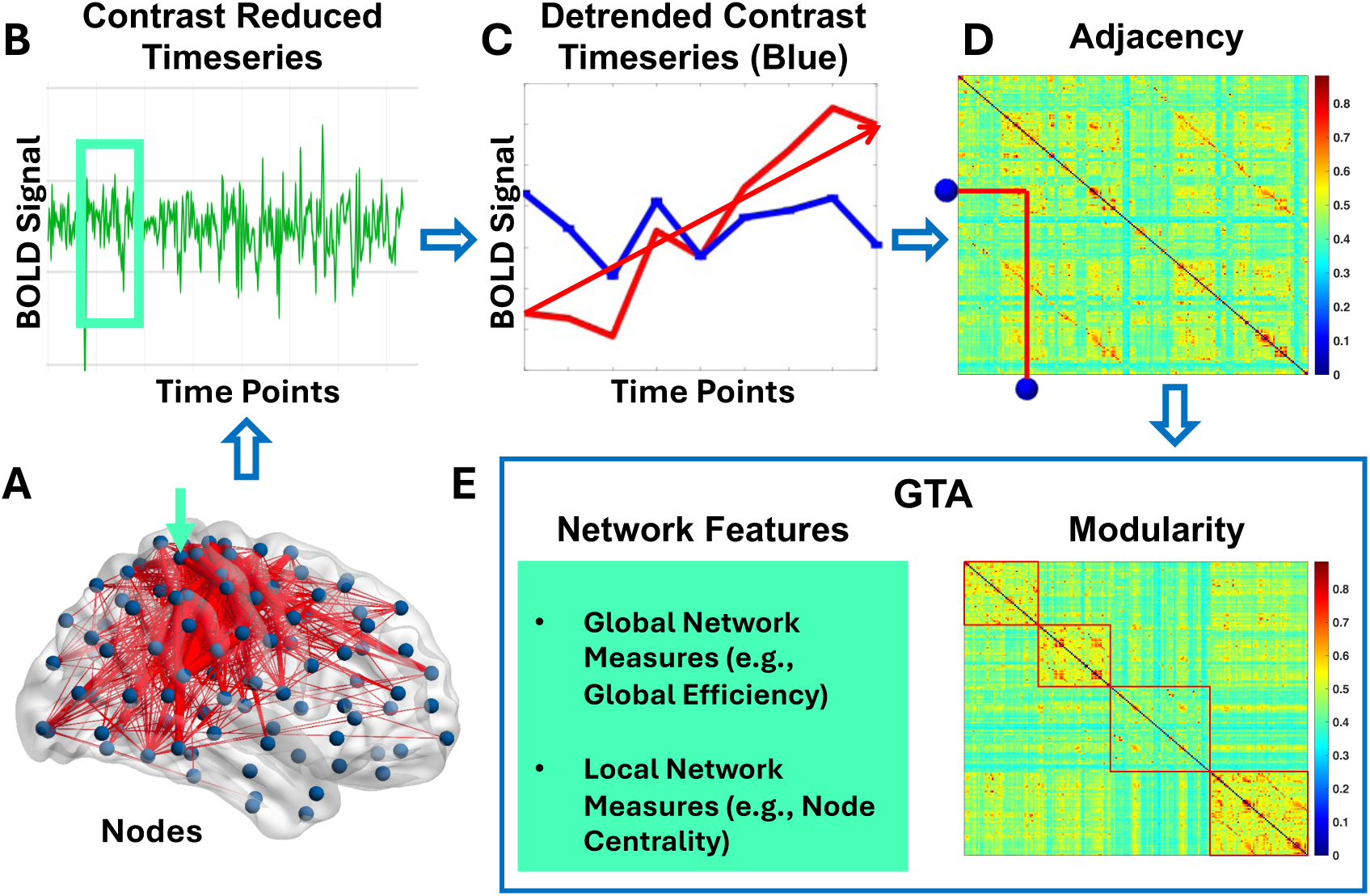
Analysis Pipeline. A, Nodes: 200 ROIs shown in blue (6mm diameter) were defined based on the centroid coordinated of the Schaefer-Yeo parcellation atlas. Red lines show example thresholded connections between them (top 5%). The cyan arrow points to the example ROI, which timeseries in the figure were averaged for. B, Contrast reduced timeseries of the example ROI an entire run: A delay predictor and GLM residuals were used to derive the contrast reduced time series. The cyan box represents a single trial (18s, 9 data points), shown in C. C, The time series from the beginning of each trial until the end of the delay period is shown in red. (the red arrow shows the linear trend) A detrending algorithm was used to derive the detrended time-series shown in blue. D, Adjacency matrices were computed for the 200 ROIs using the data with and without the trend. The example ROI indicated by the cyan arrow in A and its neighbour are shown in the x and y axis and the connection between them in the adjacency matrix is traced by the red lines. The colormap shows areas of low to high correlations (blue to red) E, GTA: The global, mesoscale and local network features were derived for each participant and trial. The modularity of the same adjacency matrix shown in D is shown by resorting the nodes based on their strongest connections and defining the different modules by the red boxes.

To provide consistent comparisons of time series across participants and conditions, for the network analysis described below, we normalized the time series from each node (e.g., Figure 2C, thick red line). This process involved creating a contrast-reduced time series for both the EGOCENTRIC and ALLOCENTRIC tasks using FSL’s FEAT(FMRIB’s Expert Analysis Tool) analysis. For each contrast vector C, the model fit F_C_ at each time point t was computed using the following formula:

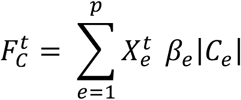

Where:

- p is the number of explanatory variables (EVs) in the design matrix,
- Xet is the value in the design matrix for EV ee at time tt,
- βe is the parameter estimate for EV ee,
- ∣Ce∣ is the absolute value of the contrast vector element CeCe.

The contrast vector C is then normalized purposes by scaling it as follows:

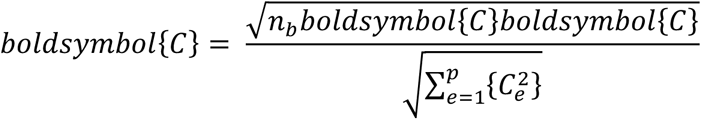

Where nboldsymbol{C} is the number of non-zero elements in the contrast vector. This ensures that partial model fits are scaled appropriately. Additionally, in the first-level analysis (i.e., fitting a model to time series data), the mean of the data is added to the model fit since FSL de-means the time series data before model fitting.

The final contrast-reduced time series was computed as the sum of the partial model fit and the residual error ɛɛ from the GLM estimation:

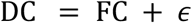

#### 2.8.1 Detrending

As in many previous sensorimotor studies (e.g., Medendorp et al., 2003; Cappadocia et al. 2017), we noted a consistent upward linear trend in the BOLD time series for most nodes (Figure 2C, thin red line). To assess the role of this general trend in network dynamics, we created a comparison dataset by linear detrending using the detrend function in MATLAB. The detrend function automatically fits a first-degree polynomial (a straight line) to the data y using a least squares approach. It then subtracts the fitted line from the data, effectively removing the linear trend and leaving only the residuals. The goal was to assess what network properties were missing after detrending, and what properties remained in the remaining fluctuations around the linear trend (Figure 2C, blue line).

#### 2.8.2 Network Analysis

Functional ‘connectivity’ edges (Figure 2A, red lines) were computed by taking the absolute values of correlations between all the nodes, computed for each participant, trial and separated into condition. Note that these edges represent the degree of temporal signal sharing between nodes, not anatomic connections. These correlations were then used to construct an ‘adjacency matrix’ for each participant / condition (Figure 2D), for both original and detrended data. Each entry of this matrix represents the correlation between nodes in the corresponding rows and columns. For example, the two blue dots illustrate the connection between the node highlighted by the cyan arrow and it’s neighbour. These matrices were then used for graph theory analysis.

### 2.9 Graph Theory Analysis (GTA)

Following the construction of the adjacency matrices, we evaluated the topological and dynamical properties of the functional brain networks for both the EGOCENTRIC and ALLOCENTRIC tasks (Figure 2E). Three common graph-theoretical measures were used to assess the network’s topology: clustering coefficient (CC), global efficiency (EF), and eigenvector centrality (EigC), as explained below (Bonacich, 1972; Latora & Marchiori, 2001; Sporns et al., 2005; Watts & Strogatz, 1998).

#### 2.9.1 Measures of Network Topology

##### 2.9.1.1 Global Network Topology

**Clustering Coefficient (CC)** measures the level of local processing within the network, indicating how interconnected the nodes are. Networks with a high CC reflect functional segregation, with nodes highly connected to one another. This is important because the brain typically organizes itself into segregated modules for efficient neural processing. In a weighted network, connectivity strength between three nodes (triangles) indicates the clustering coefficient.

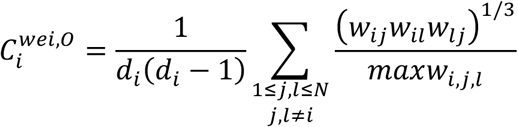

*d_i_*: The *degree* of the a given node *‘i’*
*w_ij_*_-_: The weight of connection between the pair of nodes *(‘i’* and *‘j’)*
*maxw_i,j,i_*: maximum weight between neighbor nodes that make a triangle (Saramäki et al., 2007).

**Global Efficiency (EF)** quantifies the ability of the network to integrate information across distributed regions. Higher global efficiency indicates that the network can rapidly and effectively exchange information between distant nodes, facilitating efficient brain function. In a weighted network, EF is inversely related to strongest connection between pairs nodes. This measure is average over all pairs in the network.

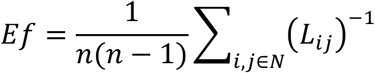

*L_ij_*: Strongest connection between pair of nodes (*i* and *j)*
*n*: Total of number of nodes (Rubinov & Sporns, 2010)

##### 2.9.1.2 Local Network Topology (Node Hubness)

**Eigenvector Centrality (EigC)** measures the influence of a node based on its connections to other well-connected nodes (Bonacich, 2007). Nodes with high eigenvector centrality are often considered hubs (Ghaderi et. al, 2023), as they are central to the flow of information and have strong connections to other highly influential nodes. However, hubs with high Eigc lack diverse connections, giving rise to their role of within-module hubs. In a weighted network, the eigenvector corresponded to largest eigenvalue of adjacency matrix is *Eig_C_*.

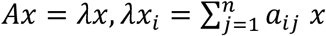

*A*: Adjacency matrix
*λ*: The largest eigenvalue of *A*
*x*: Eigenvector associated with *λ*
*n*: Total of number of nodes

**Betweenness centrality** is a measure of centrality in a graph that quantifies the extent to which a node (or edge) acts as a "bridge" between other nodes. It is based on the idea that a node is important if it lies along the shortest paths (geodesics) between other pairs of nodes in the network (Freeman, 1977). Nodes with high betweenness centrality fulfill the role of hubs (Sporns, 2007), since they are important for the flow of information, but act as between-module hubs due to their diverse connections. In other words, betweenness centrality captures how often a node is an intermediary or a connector in the network, influencing the flow of information, goods, or influence between different parts of the network. For a weighted graph, where edges have weights representing distances, costs, or capacities, the definition of the shortest path must take the weights into account. In this case, the betweenness centrality formula is adjusted to consider the weighted shortest paths between pairs of nodes.

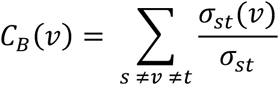

σst(v): the number of shortest weighted paths between nodes s and t that pass through node v
σst: the total number of shortest weighted paths between nodes s and t.

Significant network hubs were then determined by comparing the eigenvector centrality and betweenness centrality values across all brain regions (200 Schaefer-Yeo ROIs) using a penalized regression binary logistic model (glmnet package, R Core Team, 2024). This analysis utilized the LASSO (Least Absolute Shrinkage and Selection Operator) regression is a form of penalized regression that adds a penalty (L1 penalty: encourages sparse solutions by shrinking some coefficients to zero) to the loss function. This provides both regularization (preventing overfitting) and variable selection (shrinking some coefficients to zero). This approach compares multiple nodes at once and only leaves behind the nodes with a large difference as significant predictors.

The clustering coefficient, global efficiency, eigenvector centrality and betweenness centrality functions in the Brain Connectivity Toolbox (Rubinov & Sporns, 2010) were used to determine those measures.

#### 2.9.2 Dynamical Graph Theory Measures

Additionally, we employed graph theory analysis to investigate the dynamics of network synchronization across all pairs of brain regions (Bassett & Sporns, 2017; Breakspear, 2017; Ghaderi et al., 2020; Honey et al., 2010; Vidaurre et al., 2018). In this analysis, the following measure was used:

**Energy (H)**, which reflects the stability of synchronization in the network. Higher energy values suggest stable coupling between nodes over time, indicating more consistent functional interactions. In a weighted network, the *H* is the summation of absolute values of eigenvalues.

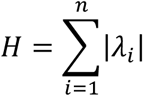

*λ_i_*: Set of network eigenvalues
*n*: Total of number of nodes (Ghaderi et al., 2020)

#### 2.9.3 Modularity

Modularity refers to a measure of the structure of a network, which quantifies the degree to which a network can be divided into relatively independent, densely connected groups or communities. A *community* in this context is a subset of nodes that are more densely connected to each other than to nodes outside the subset.

Mathematically based on Newman modularity algorithm (Newman, 2006), modularity QQ is defined as:

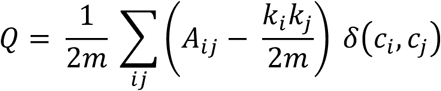

Where:

- Aij: Adjacency matrix (with 1 if there is an edge between nodes ii and jj, and 0 otherwise),
- ki: degree of node i,
- m: total number of edges in the network,
- ci: community assignment of node i,
- δ(ci,cj): Kronecker delta function, which is 1 if nodes ii and jj are in the same community, and 0 otherwise.

The modularity function in the Brain Connectivity Toolbox (Rubinov & Sporns, 2010) was used to find subnetworks. In addition, the function takes into account the resolution parameter γ which adjusts the scale of community detection, influencing the size and granularity of the modules (communities) detected in the network.

##### 2.9.3.1 Modularity and communication analysis

To evaluate modularity of subnetworks we implemented a self-developed function (Ghaderi et al., 2025) on the Newman modularity algorithm (Newman, 2006) in MATLAB 2022b. It is available as a open access code at https://github.com/AHGhaderi/Amir-HosseinGhaderi/commit/df636b2105e578a8969ffa88856e54f3267a40a7).

Steps in this algorithm are (Ghaderi et al., 2025):

1) Identify a group of nodes (e.g., brain regions) to form a subnetwork.
2) Calculate the average connectivity between the selected nodes.
3) Randomly shuffle the connections between nodes in the entire network.
4) Recalculate the average connectivity between the selected nodes after randomizing the network.
5) To find the modularity score, divide the connectivity average from step 2 (original subnetwork) by the connectivity average from step 4 (randomized network).

If the modularity score in step 5 is greater than 1, it suggests that the connectivity pattern within the selected group of nodes (subnetwork) is more structured than a random connectivity pattern. This implies that the selected nodes are connected in a modular fashion, forming a modular subnetwork.

#### 2.9.4 Null Normalization

The graph network measures, including the clustering coefficient, global efficiency and energy, were normalized using a null model generated using a self-developed Matlab code. It is available for open access at: https://github.com/AHGhaderi/Null-model.

A null model is typically used in network science to create a random version of the data, without any of the underlying structure or dependencies present in the original adjacency matrix (Váša, & Mišić, 2022). Using the Matlab function, a "null matrix" was produced for each adjacency matrix (individual trials within runs). This was done by permuting elements within the adjacency matrix in a random fashion. For each node (row), this is done by randomly shuffling elements from the lower triangular part (below the diagonal) and placing them in the upper triangular part of the matrix. The matrix is then symmetrized to maintain the structure of the original input matrix, but with randomized values. The graph network measures are then computed again from the average null matrix. The final reported values were found by dividing the value of the graph network measure in the original data by the value of the graph measure found in the average null matrix, e.g., Clustering coefficient = CC_raw matrix_ / CC_null matrix_

### 2.10 Statistical Analysis

To explore the differences between the EGOCENTRIC and ALLOCENTRIC tasks in terms of their functional brain network properties, nonparametric permutation tests were performed. These tests compared the whole network measures—such as clustering coefficient (CC), global efficiency (EF), energy (H), and Modularity (M)—between the two tasks.

The original study obtained a medium to large, standardized effect size of d = 0.67, but only analyzed 12 participants after exclusion criteria were applied. To account for the relatively small sample size (by current standards), which did not meet distributional assumptions, permutation testing was used. We randomly shuffled task conditions between participants and recalculated the t-statistic for each permutation. This process was repeated 5000 times to ensure that the resulting distribution of t-values approximated a normal distribution, allowing us to determine statistical significance at a 0.05 level. Multiple comparisons were corrected using false discovery rate (FDR) analysis to account for the potential inflation of Type I error.

The resolution limit problem arises when the resolution parameter γ is set too high or too low, causing the modularity optimization to miss communities that are smaller or larger than what the parameter was tuned for (Blondel et al., 2008; Fortunato, 2010; Good et al., 2010; Traag et al., 2011). To account for this, the modularity and number of modules was determined at different values of γ to see which gives the most meaningful community structure (including γ=1 - 1.3). A linear multilevel regression module (lme4 package, R Core Team, 2024) was used to determine task and trend differences in the number of modules at different resolution parameters, while a quadratic multilevel model was used to determine differences in modularity at different resolution parameter.

### 2.11 Classification and Predictive Modeling

We employed a machine learning approach to determine which network features (clustering coefficient, global efficiency, energy, and entropy and modularity) best differentiate between the EGOCENTRIC and ALLOCENTRIC tasks. We used a supervised cubic Support Vector Machine (SVM) classification algorithm. A cubic SVM is an extension of the SVM algorithm, where the kernel function used is the cubic kernel. This is a non-linear classification method where the decision boundary between classes is learned by transforming the input data into a higher-dimensional space, making it easier to find a linear separation in that space. In general, an SVM aims to find the best hyperplane that separates data points of different classes with the maximum margin. When the data is not linearly separable in the original feature space, we can use a kernel approach to map the data to a higher-dimensional feature space where a linear separator may exist.

To prevent overfitting, fivefold cross-validation was used during the training phase, which is a widely recognized and effective method in machine learning. This approach enhances the reliability of the results by dividing the dataset into five distinct subsets, or "folds." In each iteration, the model is trained on four of these subsets and tested on the remaining one. This process is repeated five times, with each fold serving as the test set once. By systematically using different portions of the data for training and testing, this technique minimizes the risk of overfitting and provides a more comprehensive assessment of the model’s generalizability across diverse data.

We tested different classification model’s ability to classify the EGOCENTRIC and ALLOCENTRIC conditions accurately using the Classification Toolbox in MATLAB. The MATLAB Classification Toolbox provides a variety of models for classification tasks, each suitable for different types of data and tasks, ranging from simple linear models (Logistic Regression, LDA) to complex non-linear models (SVM, Neural Networks, Random Forests). Depending on the nature of the data (e.g., linear vs. non-linear, large vs. small datasets), the best classification model is selected. The overall performance of the best model was assessed using the receiver operating characteristic (ROC) curves, which helps evaluate the model’s ability to distinguish between positive and negative cases across various thresholds by calculating the Area Under the Curve (AUC). A higher AUC indicates that the model is better at making accurate predictions, with a larger separation between the positive and negative classes, which generally reflects better overall performance. We reported the model’s overall accuracy and presented confusion matrices, including positive predictive values (PPV) and false discovery rates (FDR), to provide deeper insights into the model’s classification outcomes and reliability. The most important feature was determined using the Kruskal-Wallis importance score, which quantifies the relative contribution of each feature in explaining the variance between different groups. All classification analyses were performed in MATLAB (R2022b).

## 3. Results

In this study, we applied graph theoretical analysis to investigate network properties in a previously published fMRI dataset (Chen et al. 2024). To summarize, participants were shown a set of visual stimuli (targets and landmarks), instructed to either remember the target location and ignore the landmark (*Egocentric* task) or remember target relative to a visual landmark (*Allocentric* task), and then after a delay in which the landmark shifted, they reached toward the goal (**Figure. 1**). Univariate analysis confirmed ‘ventral stream’ coding of landmark-centered goals versus ‘dorsal stream’ coding of egocentric visual goals, with further activation in parietofrontal reach pathways during movement execution (Chen et al. 2014). Other studies suggested that ego-allocentric integration is completed in frontal cortex (Chen et al. 2018; Bharmauria et al. 2020, 2021; Schutz et al., 2023).

Based on these results, we hypothesized that the *Allocentric* condition would induce a general increase in cortical signal sharing: specifically, increased sharing of information between dorsal and ventral pathways and recruitment of additional hubs in temporal and frontal cortex. We further investigated whether removal of the general rising trend of BOLD activation would decrease network parameters related to global integration. To these ends, we examined functional network properties at the whole brain level (global network parameters), intermediate level (network modularity), and local level (network ‘hubs’), and then used a classifier model to test if the task instruction (egocentric vs. allocentric) could be decoded from these network parameters.

Note that the original study of Chen et al. (2014) reported that participants accurately pointed toward the originally viewed target location (or opposite in the anti-reach task) in the *Egocentric* Task and accurately shifted pointing direction with the landmark in the *Allocentric* Task. This suggests that participants engaged the egocentric system and effectively ignored intervening shifts in the former task but effectively utilized landmark-centred codes in the latter task. Based on this, we henceforth refer to the functional networks associated with these tasks as the egocentric and allocentric networks.

### 3.1 Whole Brain Analysis: Cortical Trends Mediate Global Network Parameters

To investigate the global topological and dynamical attributes of our networks, we computed three distinct metrics: Clustering Coefficient, Global Efficiency, and Energy (**Figure 3**), which assess network segregation, integration, and stability / synchronization of connections between nodes respectively. Each ‘violin plot’ shows the distribution of these scores across participants for the Egocentric network (blue) and allocentric network (red), contrasting scores from the original dataset (left) versus the detrended dataset (right). If overall temporal trends in brain activation mediate long range communications, detrending should result in increased local clustering and decreased integration. We found this to be the case for two of our parameters: detrending the data increased network segregation, as indicated by a significant rise (0.048 increase, t(df=22) = 3.64, p= 0.0034) in the average clustering coefficient of nodes in both tasks (**Figure 3A**) and significantly reduced synchrony (**Figure 3C**) when the data were detrended (0.386 decrease, t(df=22) = 5.20, p= 2e-04). However, there was no significant effect on Global Efficiency (**Figure 3B**) and none of the measures were significantly modulated by task. To discriminate the influence of behavior on these networks, we next examine the intermediate level: cortical modularity in both tasks, in original and detrended datasets.

**Figure 3:**
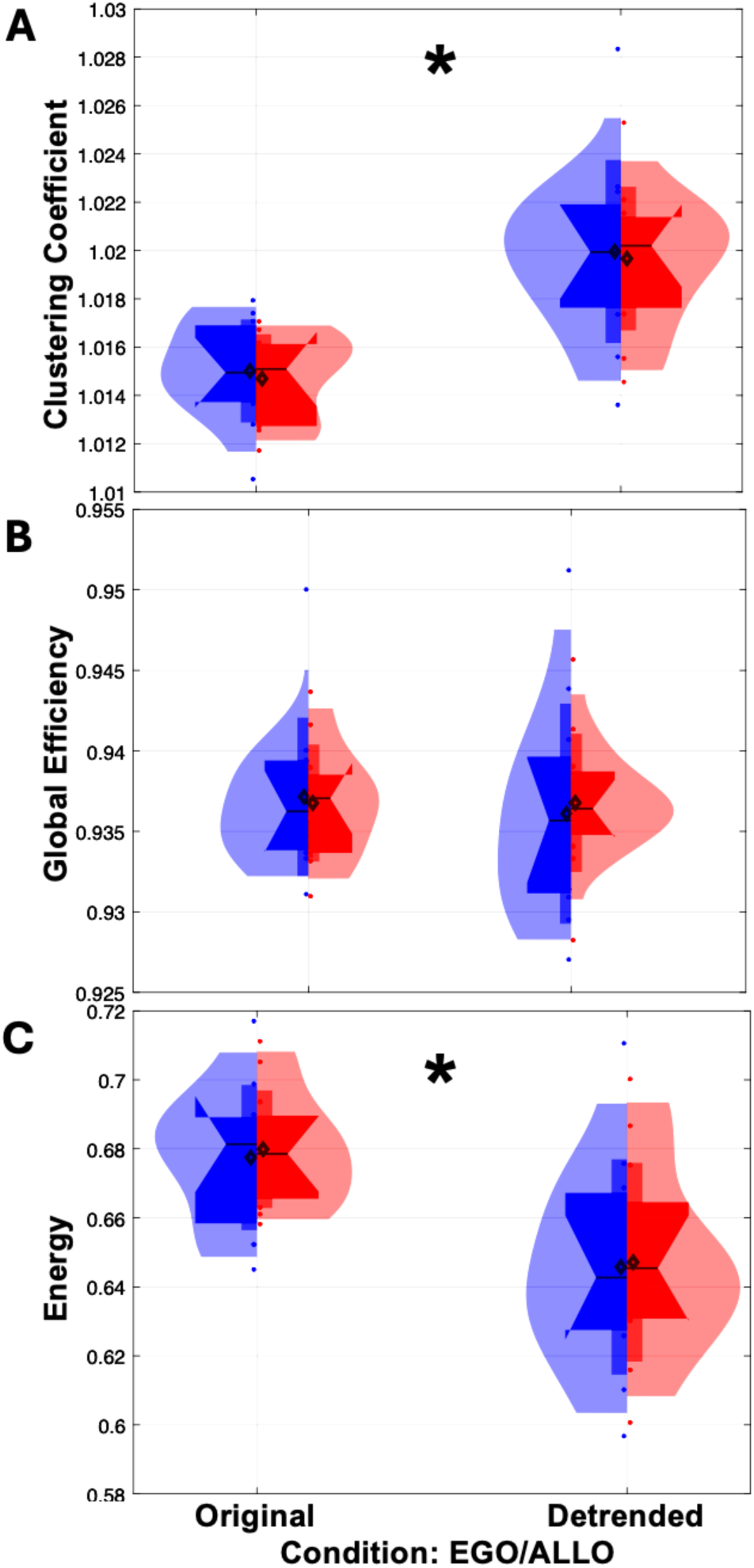
Quantification of Global Network Properties. The average clustering coefficient (**A**), global efficiency (**B**) and energy (**C**) for each participant (n=12) were compared between the Egocentric (blue) and Allocentric networks, for the original (left) and detrended (right) data.

### 3.2 Intermediate Level Analysis: Behavior Influences Signal Modularity

#### Graphic Analysis

Modularity analysis sorts nodes into ‘communities’ with signals that tend to correlate with each other and not to the other communities. **Figure 4** provides a graphic summary of our general network and modularity analysis for group data from the *Egocentric* condition (A), *Allocentric* (B), and detrended *Allocentric* data (C). The small dots indicate the location of the nodes analyzed, the larger dots local network hubs (these will be discussed in a later section) and the lines indicate ‘edges’ (signal correlations) that exceeded the 90^th^ percentile correlation (thin lines) or 95^th^ percentile correlation (thick lines). Note that the nodes, hubs, and edges have been color-coded based on the results of a modularity analysis at standard (1.0) resolution parameter γ.

**Figure 4:**
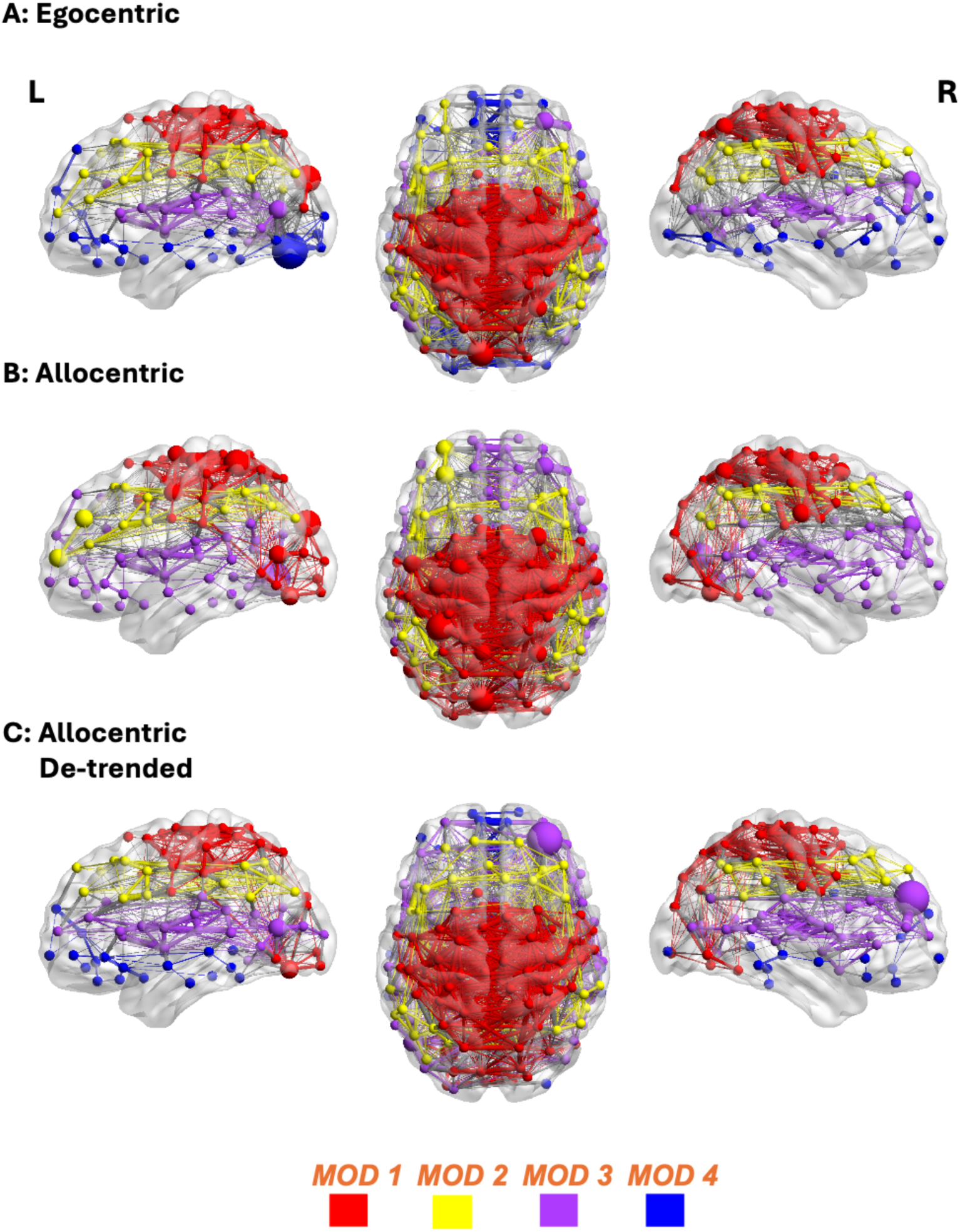
**Visualization of Networks / Modularity Analysis**. *Egocentric* (**A**), *Allocentric* **(B**) and *detrended Allocentric* (**C**) datasets are shown. Each row provides an ‘above’ view of bilateral cortex along with lateral views of the left and right hemispheres. Small dots: node locations. Only nodes that are connected to edges exceeding the threshold shown below are shown. Larger dots: Eigenvector network hubs (scaled by statistical significance-based importance). Lines: ‘edges’ (signal correlations) exceeding a threshold of 90^th^ percentile correlation (thin lines) or 95^th^ percentile correlation (thick lines). Nodes, hubs, and edges have been color-coded by module (up to four): yellow, purple, blue, and red. For this analysis a standard modularity resolution parameter of 1.0 was used.

As shown in **Figure 4 A**, the *Egocentric* network was divided into four bilateral modules: a superior occipital-parietal / somatomotor module (Module 1; red).an inferior parietal / lateral frontal module (Module 2; yellow), a superior temporal / inferior frontal module (Module 3; purple), an inferior occipital-temporal / prefrontal module (Module 4; blue), and In contrast, the *Allocentric* network (**Figure 4 B**) was divided into three modules, because the blue module 4 was incorporated into modules 1 and 3, forming an inferior occipital / superior parietal / somatomotor module (red), and a temporal / inferior frontal module (purple). Only the inferior parietal / lateral frontal module (2, yellow) remained the same.

In light of our hypotheses, it is noteworthy that the *Allocentric* network showed an increased conjoining of ventral and dorsal stream nodes (compared to Egocentric), toward a single (red) occipital / superior parietal / somatomotor module. However, after detrending the data (Figure 4C), this network reverted to four modules and the conjoining of dorsal-ventral visual stream modules was reduced. We will quantify these and other observations in the following sections.

#### Quantitative Analysis

F**igure 5** shows the distributions of modularity score across participants for each module (color coded red, yellow, purple, blue). Each color plot is further segregated into scores for the egocentric (left halves) and allocentric (right halves) networks. In simple terms, higher distributions signify greater modularity: increased connectivity within the module and reduced connectivity without.

**Figure 5:**
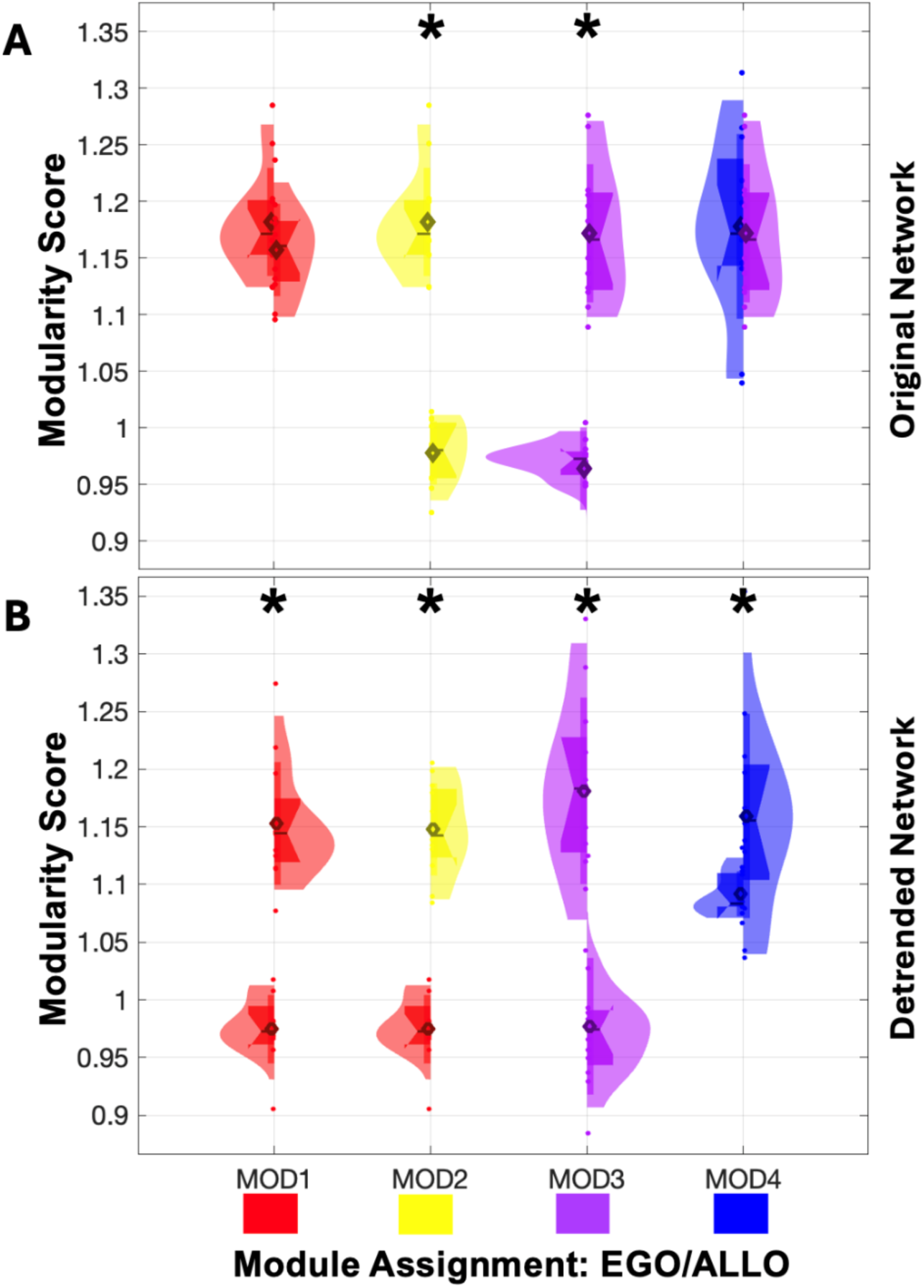
Quantification of Modularity in Original and Detrended Datasets. The left halve of each violin plot shows Egocentric network data and the right half shows Allocentric network data. A standard resolution parameter of 1 is used here. The color key is based on the original modules shown in Fig. 4. **A**: Original Ego/Allocentric datasets. Since Egocentric module 4 (blue) disappeared in the Allocentric dataset, mostly joining module 3, it is compared to allocentric module 3 (purple). **B**: Detrended Ego/Allocentric datasets. Since Allocentric modules 1 and 4 were merged in the detrended Egocentric dataset, the same values (red) were used for comparison to the Allocentric modules. Significant differences between task are indicated with an asterisk ‘*’ in the plot.

**Figure 5 A** quantifies modularity in the original dataset. Most of the original modules, with the exception of Allocentric module 2 (yellow) and Egocentric module 3 (purple), had a connectivity pattern that was more specific than a random null distribution. Comparing between tasks, the *Egocentric* network exhibited significantly higher modularity in the inferior-parietal / lateral frontal module (yellow) (1.25 increase, t(df=22) = 4.43, p= 8e-04), whereas the *Allocentric* Network demonstrated significantly higher modularity in the superior temporal / inferior frontal module (purple) (1.29 increase, t(df=22) = 4.00, p= 0.00). This suggests a trend toward greater dorsal modularity in the egocentric task vs. ventral modularity in the Allocentric task, consistent with dorsal-ventral stream theory.

As noted above (**Figure 4 C)**, the Allocentric network reverted to four modules after detrending **(Figure 5 B**). Detrending produced dramatic changes in the distribution of modularity coefficients. In this case, only Egocentric module 3 had a connectivity pattern more specific than a random null distribution, whereas Allocentric modules 1, 2, 3 were no more specific relative to a random null distributions or the corresponding egocentric modules. Together with **Figure 4**, this analysis suggests that the trend plays a role in integrating ventral occipital and dorsal parietal regions in the Allocentric network, as well as in forming specialized modules within the dorsal stream.

#### Consistency Across Resolution Parameters

The proceeding modularity analysis was performed with and arbitrary standard resolution parameter of 1.0, but GTA parameters are known to be parameter-dependent (Blondel et al., 2008; Fortunato, 2010; Good et al., 2010; Traag et al., 2011). Therefore, it is possible that we just ‘got lucky’ by using the standard parameter. To assess the influence of this arbitrary choice on task- and trend-dependence, we calculated modularity scores and the number of modules for the original and detrended datasets across a range of resolution parameters (**Figure 6**).

**Figure 6.**
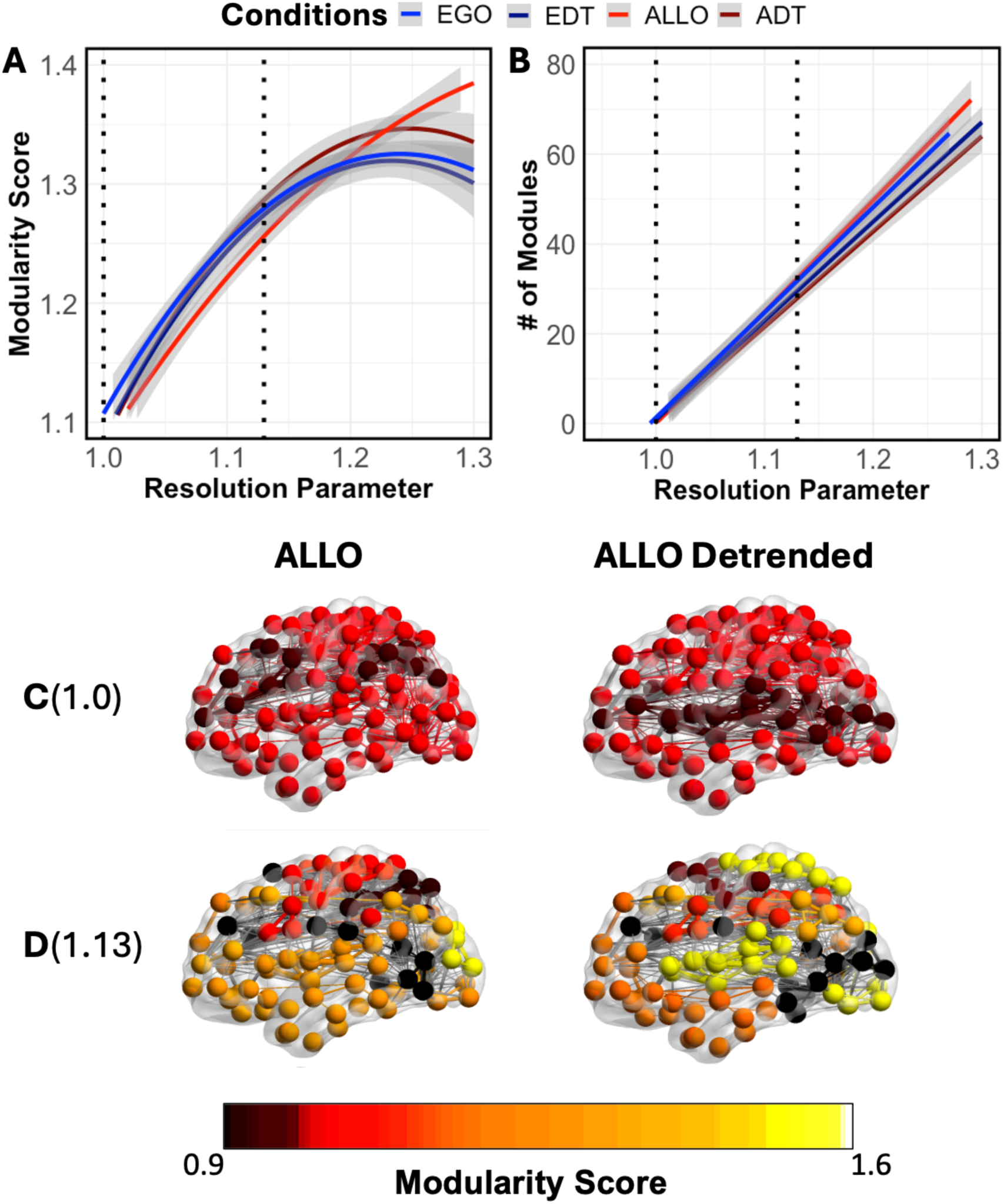
Influence of Threshold and Detrending. The change in the modularity score (**A**) and the number of modules (**B**) is shown as a function of resolution parameter (top) is shown in the Egocentric (blue) and Allocentric (red) task, in the original (EGO/ALLO) and detrended (EDT/ATD) networks (darker colors). Network connectivity is shown in different resolution parameters for the Allocentric dataser, **C**: 1.0 (Standard) and **D**: 1.13 (Optimal for ALLO) (bottom), for original (left) and detrended (right data). The nodes are separated based on the module assignments based on the respective resolution parameters but color coded based on the modularity score from part **B**.

Across different resolution parameters, we found that the influence of task was best revealed in the modularity score (**Figure 6 A**). The relationship between resolution and mean modularity was quadratic (e.g. for Egocentric modularity at a resolution parameter of 1, mean modularity score = average modularity score of all 4 egocentric modules in Figure 5). All scores generally rose with resolution, with original *Egocentric* data (blue) showing significantly higher scores (0.137, t(596) = 1.69, p =0.011) than original *Allocentric* (red) below 1.18, and significantly lower above 1.18 resolution (-0.161, t(596) = -1.79, p =0.009). The detrended data (dark red / black) essentially followed the same curves. It is noteworthy that both the standard (1.0) and optimal (1.13) resolutions (vertical dotted lines) fall to the left of this reversal point. Color coded modularity scores of individual nodes for the *Allocentric* networks are illustrated in the graphic plots shown below, where the score for most nodes rises (lighter colors) from standard resolution (**C**) to optimal resolution 1.13 (**D**) and show slightly different topological patterns for original (left) vs. detrended (right) data, such as a reduction in occipital modularity. In contrast, the influence of resolution parameter on module number was linear (**Figure 6 B**), and not task dependent. Instead, it showed significantly different slopes 0.31 difference (t(596) = 1.98, p = 0.003) for original (red/blue) vs. detrended (dark red/black) data. Overall, **Figure 6** suggests that, although modularity depends on the resolution parameter, task- and trend-dependent differences are retained across different values.

### 3.3 Local Analysis: Hubness of Egocentric and Allocentric Brain Networks

Hubs are nodes that highly correlate to other areas that in turn are well ‘connected’ to other nodes and are thus thought to play important roles in the network. *Eigenvector Centrality* is used to compute locally important hubs (i.e., functional connectivity within modules) whereas *Betweenness Centrality* is used to compute hubs for global (i.e. across module) connectivity (Freeman, 1977; Rubinov & Sporns, 2010).

#### Local Hubs

**Figure 7** illustrates significant Eigenvector Centrality hubs for the significant Egocentric (**A**) and Allocentric (**B**) modules, overlaid over clusters of significant BOLD activation (orange). Eigenvector Centrality identifies nodes that are well-connected to other influential nodes (Freeman, 1977). In the figure, these nodes are scaled by size and color related to their hub importance and modularity assignment, respectively, which indicates their relative contribution to the network.

**Figure 7:**
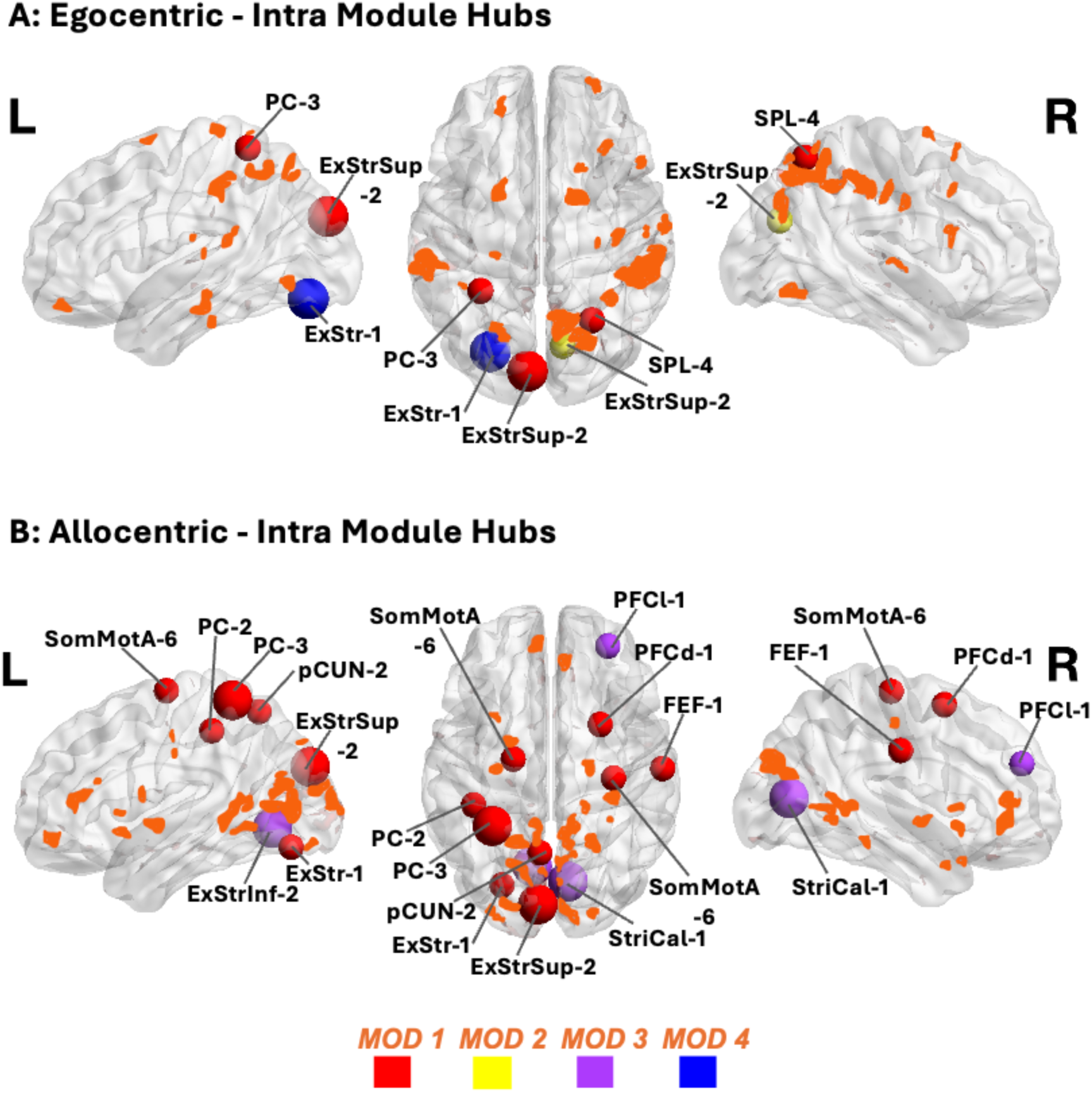
Eigenvector Centrality Hubs. **A**: Egocentric, **B**: Allocentric hubs with significant eigenvector centrality are shown overlaid on the cluster map for the Allocentric vs Baeline Mixed-effects analysis, clusters with a corrected p-value threshold of 0.0167 are shown. See Table 1 for definitions of acronyms, their coordinates, and correspondence to functional areas. The node sizes are scaled according to significance-based importance (p<0.01 larger nodes, p<0.05 smaller nodes), whereas the color scheme shown at the bottom of the figure is based on the same modularity conventions of Figures 4 & 5. Note nodes that belonged to modules with insignificant modularity are excluded, including nodes from the Module 3 in the *Egocentric* task, and nodes from Module 2 in the Allocentric task.

**Table 1:** Eigenvector Centrality Hubs of the Egocentric and Allocentric Networks.

| *ROI Name* | *Network* | *MNI-x* | *MNI-y* | *MNI-z* | *EGO EVC* |
| --- | --- | --- | --- | --- | --- |
| *LH_ExStr_1* | VisCent | -26 | -77 | -14 | 2 |
| *LH_ExStrSup_2* | VisPeri | -7 | -87 | 28 | 2 |
| *LH_PC_3* | DorsAttnB | -31 | -46 | 63 | 1 |
| *RH_ExStrSup_2* | VisPeri | 11 | -74 | 26 | 1 |
| *RH_SPL_4* | DorsAttnA | 26 | -61 | 58 | 1 |

| *ROI Name* | *Network* | *MNI-x* | *MNI-y* | *MNI-z* | *ALLO EVC* |
| --- | --- | --- | --- | --- | --- |
| *LH_ExStr_1* | VisCent | -26 | -77 | -14 | 1 |
| *LH_ExStrInf_2* | VisPeri | -10 | -67 | -4 | 2 |
| *LH_ExStrSup_2* | VisPeri | -7 | -87 | 28 | 2 |
| *LH_PC_2* | DorsAttnB | -41 | -35 | 47 | 1 |
| *LH_PC_3* | DorsAttnB | -31 | -46 | 63 | 2 |
| *LH_pCun_2* | ContC | -6 | -60 | 57 | 1 |
| *LH_SomMotA_6* | SomMotA | -20 | -11 | 68 | 1 |
| *RH_FEF_1* | DorsAttnB | 59 | -16 | 34 | 1 |
| *RH_PFCd_1* | ContA | 26 | 7 | 58 | 1 |
| *RH_PFCI_1* | SalVentAttnB | 30 | 48 | 27 | 1 |
| *RH_SomMotA_6* | SomMotA | 33 | -21 | 65 | 1 |
| *RH_StriCal_1* | VisPeri | 9 | -75 | 9 | 2 |

*Note:* EVC: Eigenvector Centrality; LH\_ExStr\_1: Left Hemisphere Extrastriate 1; LH\_ExStrInf\_2: Left Hemisphere Extrastriate Inferior 2; LH\_ExStrSup\_2: Left Hemisphere Extrastriate Superior 2; LH\_PC\_2: Left Hemisphere Parietal Cortex 2; LH\_PC\_3: Left Hemisphere Parietal Cortex 3; LH\_pCun\_2: Left Hemisphere Precuneus 2; LH\_SomMotA\_6: Left Hemisphere Somatomotor Area 6; RH\_FEF\_1: Right Hemisphere Frontal Eye Fields 1; RH\_PFCd\_1: Right Hemisphere Prefrontal Cortex Dorsal 1; RH\_PFCI\_1: Right Hemisphere Prefrontal Cortex Lateral 1; RH\_SomMotA\_6: Right Hemisphere Somatomotor Area 6; RH\_StriCal\_1: Right Hemisphere Striatum Calcarine 1; VisCent: Visual Central; VisPeri: Visual Periphery; DorsAttnB: Dorsal Attention B; ContC: Control C; SomMotA: Somatomotor A ContA: Control A; SalVentAttnB: Salience Ventral Attention B. A value of 2 in the last column indicates $p < 0.01$ and value of 1 indicates $p < 0.05$ .

In the *Egocentric* network (**Figure 7 A**) local hubs were primarily confined to posterior parietal and occipital areas, including the superior parietal lobule (SPL), post-central cortex (PC), and the inferior and superior extra striate visual cortex (ExStr, ExStrSup). Based on their coordinates (**Table 1**), Some, but not all of these hubs overlap with significant BOLD activation clusters in occipital and parietal cortex (although one cannot conclude absence of activity in the other hubs). In short, the Egocentric task engaged local hubs in areas involved associated with visual processes, attention, and the bottom-up visuomotor transformations for this task (Cappadocia et al. 2017; Blohm et al., 2019).

The *Allocentric* task (**Figure 7 B**) engaged more occipital hubs in the inferior and superior extra striate visual cortex (ExStr, ExStrInf, ExStrSup), and also engaged several dorsal hubs such as somatomotor and postcentral visual cortex areas (SomMotA, PC, Precuneus) and more anterior hubs, including the lateral and dorsal prefrontal cortex (PFCl & PFCd) and the frontal eye fields FEF) (**Table 1**). Again, some of these hubs overlap with significant BOLD activation clusters, especially in occipital cortex.

#### Global Hubs

**Figure 8** depicts significant betweenness centrality hubs within all of the Egocentric (**A**) and Allocentric (**B**) modules. Betweenness centrality is a critical measure in graph theory, representing the nodes that serve as key connectors between distinct modules, thereby facilitating information flow across the network. In the Egocentric network, hubs with the highest betweenness centrality were relatively fewer and more widely distributed across different modules.

**Figure 8:**
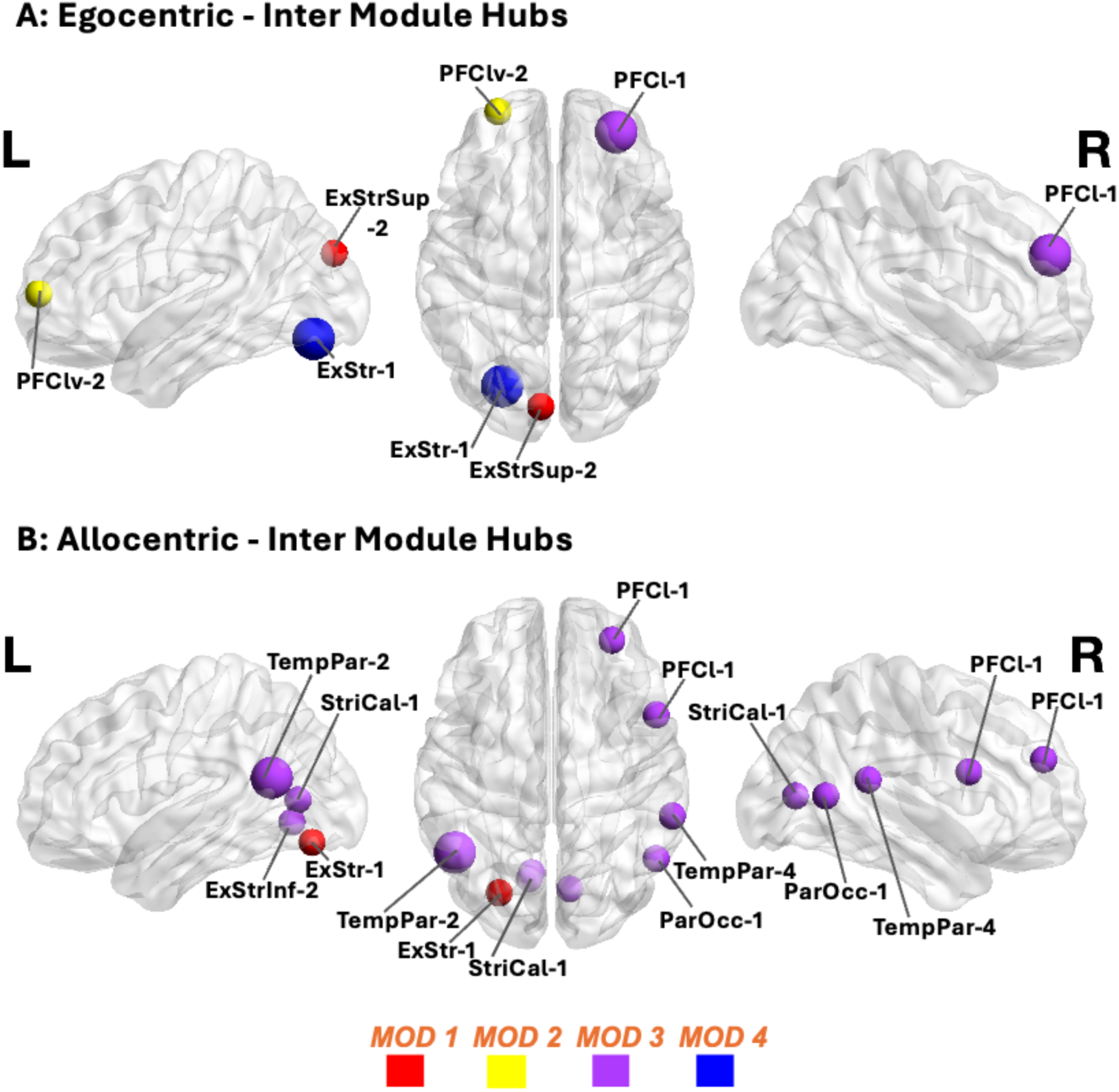
Betweenness Centrality Hubs. **A**: Egocentric **B:** Allocentric hubs with significant betweenness centrality are shown. See Table 2 for definitions of acronyms, their coordinates, and correspondence to functional areas. The node sizes are scaled according to significance-based importance (p<0.01 larger nodes, p<0.05 smaller nodes), whereas the color scheme is based on the same modularity conventions of Figures 4 & 5. Nodes from modules with insignificant modularity were not excluded from analysis, as their connectivity role could still contribute to network integration.

**Table 2:** Betweenness Centrality Hubs of the Egocentric and Allocentric Networks.

| *ROI Name* | *Network* | *MNI-x* | *MNI-y* | *MNI-z* | *EGO BTW* |
| --- | --- | --- | --- | --- | --- |
| *LH_ExStr_1* | VisCent | -26 | -77 | -14 | 2 |
| *LH_ExStrSup_2* | VisPeri | -7 | -87 | 28 | 1 |
| *LH_PFCIv_2* | ContB | -28 | 58 | 8 | 1 |
| *RH_PFCI_1* | SalVentAttnB | 30 | 48 | 27 | 2 |
| *ROI Name* | *Network* | *MNI-x* | *MNI-y* | *MNI-z* | *ALLO BTW* |
| *LH_ExStr_1* | VisCent | -26 | -77 | -14 | 1 |
| *LH_ExStrInf_2* | VisPeri | -10 | -67 | -4 | 1 |
| *LH_StriCal_1* | VisPeri | -11 | -70 | 7 | 1 |
| *LH_TempPar_2* | TempPar | -48 | -57 | 18 | 2 |
| *RH_ParOcc_1* | DorsAttnA | 52 | -60 | 9 | 1 |
| *RH_PFCI_1* | SalVentAttnB | 30 | 48 | 27 | 1 |
| *RH_PFCI_1* | ContA | 52 | 11 | 21 | 1 |
| *RH_StriCal_1* | VisPeri | 9 | -75 | 9 | 1 |
| *RH_TempPar_4* | TempPar | 60 | -39 | 17 | 1 |

Note: BTW: Betweenness Centrality; LH\_PFCIv\_2: Left Hemisphere Prefrontal Cortex Lateral Ventral 2; RH\_PFCI\_1: Right Hemisphere Prefrontal Cortex Lateral 1; LH\_TempPar\_2: Left Hemisphere Temporal Parietal 2; RH\_ParOcc\_1: Right Hemisphere Parietal Occipital 1; RH\_TempPar\_4: Right Hemisphere Temporal Parietal 4; TempPar: Temporal Parietal; DorsAttnA: Dorsal Attention A. A value of 2 in the last column indicates $p < 0.01$ and value of 1 indicates $p < 0.05$ .

In contrast, the Allocentric network exhibited additional betweenness centrality hubs, predominantly within the temporal (purple) module, with the notable exception of the extrastriate visual cortex (ExStr), part of the inferior occipital / superior parietal / somatomotor (red) module. The Allocentric hubs within the temporal module were concentrated in the ventral occipital cortex, including the inferior extrastriate cortex (ExInf) and the striate calcarine (StriCal), as well as in the temporoparietal (TempPar) and paraoccipital visual cortices (ParOcc), and extended to the prefrontal cortex (Table 2). In short, it appears that additional hubs were recruited for both local and global functional connectivity.

#### Allocentric vs. Egocentric Hubs

To directly test the influence of task instruction, we compared hubness in all nodes of the allocentric vs. egocentric networks. (Note that nodes that showed significantly different ‘hubness’ were not necessarily hubs in either individual network). **Figure 9** illustrates nodes with significantly increased Eigenvector Centrality (**A**) and Betweenness Centrality (**B**) in the Allocentric network compared to the Egocentric network. Areas that showed significantly higher Eigenvector centrality in the Allocentric task included the left orbitofrontal cortex (OFC), left parietal operculum (ParOper), and left dorsolateral prefrontal cortex (PFCd), as well as regions in the right hemisphere like the insula (Ins), inferior parietal lobule (IPL), and somatomotor areas (SomMotA_1, SomMotA_4, SomMotA_9). Note that these were distributed across both the ‘red’ occipital / superior parietal / somatomotor module 1 and the ‘purple’ temporal module 3. In contrast, only two regions showed significantly increased Betweenness Centrality in the Allocentric task: inferior extrastriate cortex (ExStrInf) and the left temporoparietal junction (TempPar), and these were confined to module 3. Overall, the allocentric network was associated with widespread increases in hubness across all four lobes of cortex.

**Figure 9:**
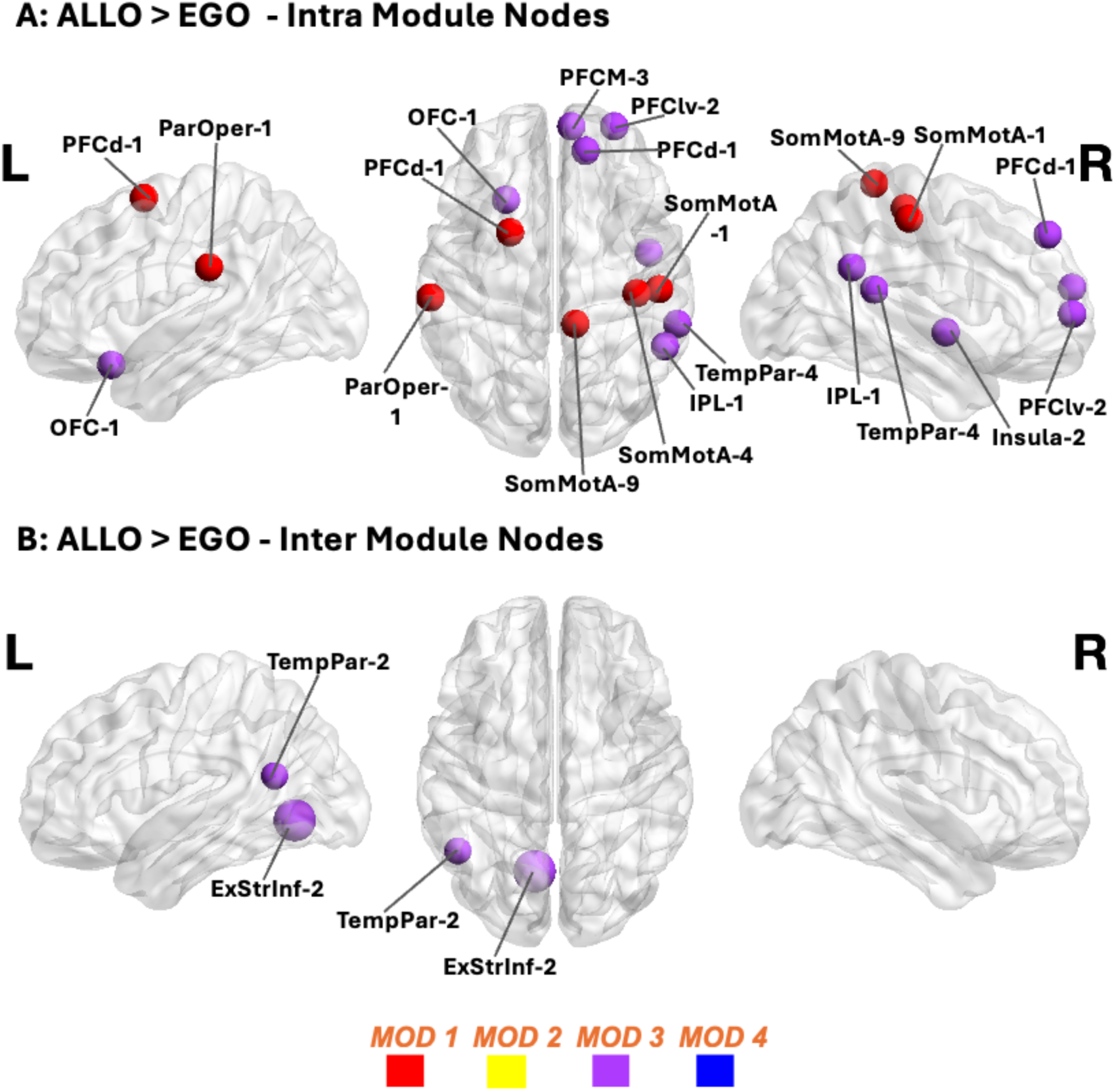
Allocentric Inter-task Difference Nodes. A: Eigenvector Centrality B: Betweenness Centrality nodes with significantly higher values in the Allocentric task (ALLO) than the Egocentric task (EGO) are shown. See Table 3 for definitions of acronyms, their coordinates, and correspondence to functional areas. The node sizes are scaled according to significance-based importance (p<0.01 larger nodes, p<0.05 smaller nodes), whereas the color scheme is based on the same modularity conventions of Figures 4 & 5.

**Table 3:** Task Difference Allocentric Nodes - Higher Eigenvector Centrality and Betweenness Centrality than Egocentric Network.

| *ROI Name* | *Network* | *MNI-x* | *MNI-y* | *MNI-z* | *ALLO*<br>*EVC* |
| --- | --- | --- | --- | --- | --- |
| *LH_OFC_1* | LimbicB | -24 | 22 | -20 | 1 |
| *LH_ParOper_1* | SalVentAttnA | -61 | -26 | 28 | 1 |
| *LH_PFCd_1* | ContA | -22 | 6 | 62 | 1 |
| *RH_Ins_2* | SalVentAttnA | 46 | -4 | -4 | 1 |
| *RH_IPL_1* | DefaultA | 54 | -50 | 28 | 1 |
| *RH_PFCd_1* | DefaultB | 15 | 46 | 44 | 1 |
| *RH_PFClv_2* | ContB | 29 | 58 | 5 | 1 |
| *RH_PFCM_3* | DefaultA | 8 | 58 | 18 | 1 |
| *RH_SomMotA_1* | SomMotA | 51 | -22 | 52 | 1 |
| *RH_SomMotA_4* | SomMotA | 40 | -24 | 57 | 1 |
| *RH_SomMotA_9* | SomMotA | 10 | -39 | 69 | 1 |
| *RH_TempPar_4* | TempPar | 60 | -39 | 17 | 1 |
| *ROI Name* | *Network* | *MNI-x* | *MNI-y* | *MNI-z* | *ALLO*<br>*BTW* |
| *LH_ExStrInf_2* | VisPeri | -10 | -67 | -4 | 2 |
| *LH_TempPar_2* | TempPar | -48 | -57 | 18 | 1 |

Note: LH\_OFC\_1: Left Hemisphere Orbitofrontal Cortex 1; LH\_ParOper\_1: Left Hemisphere Parietal Operculum 1; LH\_PFCd\_1: Left Hemisphere Prefrontal Cortex Dorsal 1; RH\_Ins\_2: Right Hemisphere Insula 2; RH\_IPL\_1: Right Hemisphere Inferior Parietal Lobule 1; RH\_PFClv\_2: Right Hemisphere Prefrontal Cortex Lateral Ventral 2; RH\_PFCM\_3: Right Hemisphere Prefrontal Cortex Medial 3; RH\_SomMotA\_1: Right Hemisphere Somatomotor Area 1; RH\_SomMotA\_4: Right Hemisphere Somatomotor Area 4; RH\_SomMotA\_9: Right Hemisphere Somatomotor Area 9. A value of 2 in the last column indicates $p < 0.01$ and value of 1 indicates $p < 0.05$ .

### 3.4 Decoding Behavior from Network Parameters

Finally, to confirm that our networks represent and *predict* meaningful behavior, and determine which parameters were most important, we decoded task (Egocentric vs. Allocentric) pooling all individual trial data across runs and participants. The decoding was done using a cubic support vector machine (SVM) classification algorithm, which was the model with the highest level of accuracy (see methods for details) and 5-fold cross-validation to evaluate the model’s performance, which provides a more comprehensive assessment of the model’s generalizability across diverse data. Our previous analysis suggested that modularity discriminates task (**Figure 4-6**), whereas global parameters did not (**Figure 3**). To test this formally, the following parameters were used initially as input to the SVM model: clustering coefficient, energy, global efficiency and modularity of the individual modules. For modules that differed between tasks, the module with the highest overlap across tasks, i.e., most shared ROIs, was used to compute modularity. This resulted in 3 modularity parameters for both the original and detrended data.

Overall, this resulted in a high level of accuracy (85.7%) in distinguishing the Egocentric and Allocentric Networks in the original network, **Figure 9** (Note that this accuracy was achieved after removing the two least important parameters, clustering coefficient and energy, based on Kruskal-Wallis feature selection method). In **Figure 9 A** we presented confusion matrices, including the positive predictive values (PPV) and false discovery rates (FDR), to provide deeper insights into the model’s classification outcomes and reliability. The model achieved a PPV of 87.2% in the *Egocentric* network, with an FDR of 12.8% and a PPV of 84.3% in the *Allocentric* network, with an FDR of 15.7%. Based on this outcome, SVM performance was similar but slightly higher for *Egocentric* network than the *Allocentric* network, the combination of modularity parameters and efficiency values have better prediction of the *Egocentric* network when using all the data.

In the *detrended* network the accuracy rose to 87.7% in distinguishing the *Egocentric* and *Allocentric* detrended networks (**Supplementary Figure 1**). The model achieved a PPV of 85.1% in the *Egocentric* network, with an FDR of 14.9% and a PPV of 90.8% in the *Allocentric* network, with an FDR of 9.2%. (**Supplementary Figure 1 A**). This means that SVM performance improves in the detrended data due to better classification of the *Allocentric* network using the combination of modularity parameters and efficiency.

**Figure10 B** shows the relative importance the features for classifying *Egocentric* versus *Allocentric* networks in the original dataset, as determined by the Kruskal-Wallis (KW) approach. Modularity was the most important parameter in classifying the two networks (*Egocentric, Allocentric*), specifically the modularity of the inferior-parietal / lateral frontal module (yellow module 2 in **Figure 4**) with a KW score of 138.72 and the modularity values of superior temporal / inferior frontal module (purple module 3 in **Figure 4**) with a KW score of 107. 85. The superior-occipital-parietal / somatomotor module (red module 1 in **Figure 4**) still had some relevant importance (3.00), but efficiency had very little importance in classification (0.73). In contrast, in the detrended network (**Supplementary, Figure 1 B**), the superior occipital-parietal / somatomotor module and the superior temporal / inferior frontal module (p) were the most important (KW of 126.51 and 92.08 respectively), but the remaining two predictors still had a high level of importance, including the occipital-temporal / prefrontal module (b) with a KW of 17.84 and efficiency with a KW of 6.31.

Finally, **Figure 10 C** uses scatter plots to visualize the distribution of data points for the Egocentric (blue) and Allocentric (orange) networks, and to qualitatively assess how well the model separated these classes. Each point provides the two most important features derived from **Figure 10 B**, plotting the modularity values of inferior-parietal-lateral frontal module 2 vs the modularity values of superior temporal / inferior frontal module. Correct classifications are shown as dots, whereas incorrect classification are shown as crosses. The *Egocentric* and *Allocentric* form clusters that appear to separate along a diagonal axis with a negative slope, with the Allocentric datapoints clustered at the upper left corner of the scatterplot, which is characterized by higher modularity in the superior temporal / inferior frontal module (y-axis) and lower modularity in the inferior parietal / lateral frontal module 2, with little overlap. The majority of the classifications were correct in the egocentric (266 correct vs. 39 mistakes) and Allocentric (279 correct vs. 52 mistakes). Overall, this analysis confirms that 1) our GTA parameters have both descriptive and predictive value for the associated behavior, and 2) that modularity had the biggest influence in discriminating the two different tasks in this dataset.

**Figure 10:**
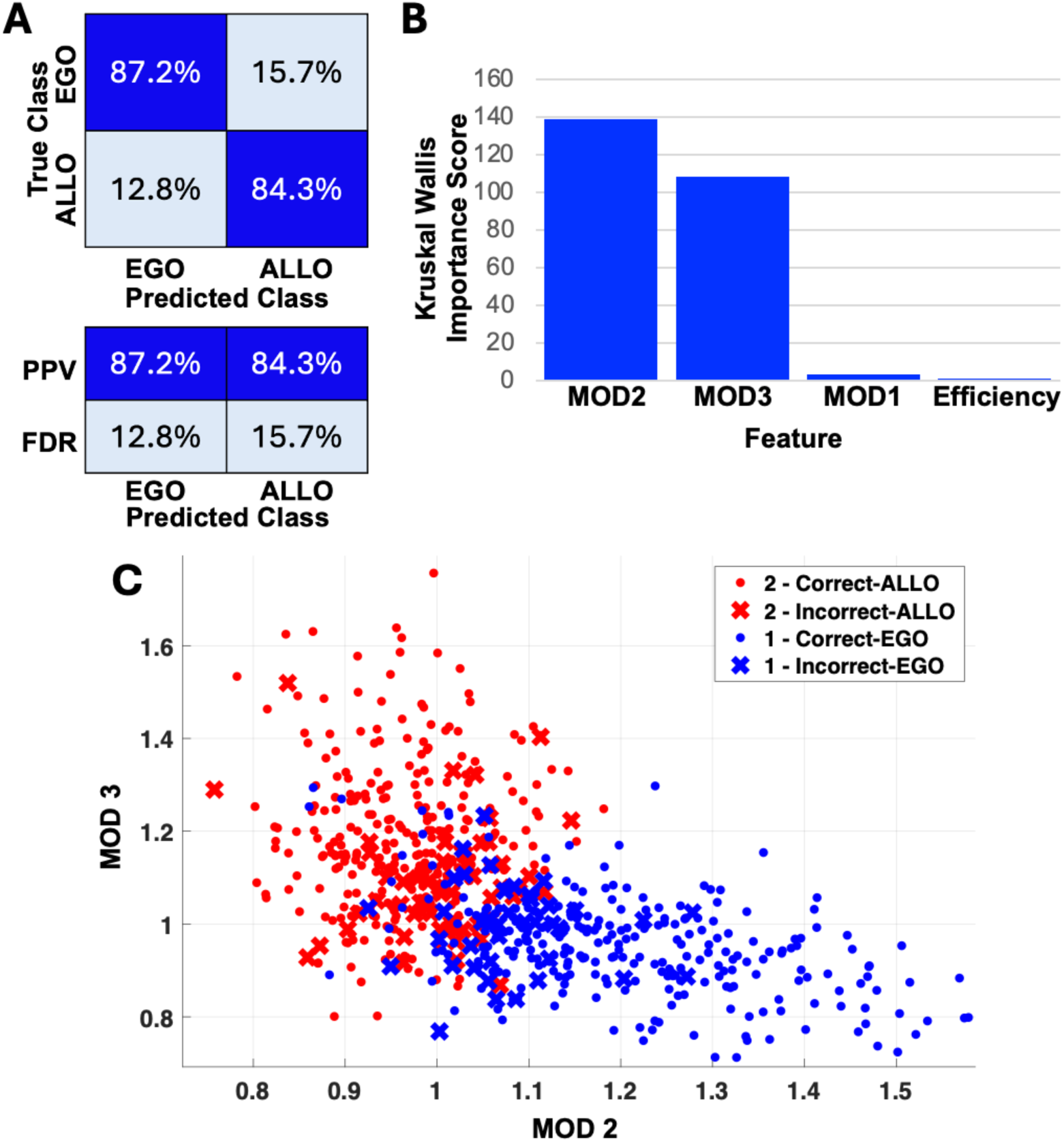
Classification Analysis. Performance of the SVM classifier in distinguishing between the Egocentric and Allocentric networks in the original network (**A**) The confusion matrix (top panel table) comparing the percentage of data points in the four categories shown, top row: data points with a true and predicted class of *Egocentric* (EGO (top left cell), data points with a true class of *Egocentric* (EGO) and predicted class of *Allocentric (ALLO)* (top left cell), and the corresponding categories for the Allocentric task in the second row. The bottom panel table shows the PPV and FDR of the *Egocentric* network (EGO, left column) and the *Allocentric* network (ALLO, right column). (**B**) Importance scores of the features classifying Egocentric and Allocentric networks in the original dataset, as determined by the Kruskal-Wallis approach. (**C**) Scatter plot of distribution of datapoints predicted across two dimensions: (modularity score of module 2 [MOD 2] vs modularity score of module 3 [MOD 3) out of the total four classifier dimensions. Correct classifications are shown as dots whereas misclassifications are shown as crosses. The two networks being classified, *Egocentric and* Allocentric are shown in blue and red, respectively.

## 4. Discussion

The present study examined the influence global activation trends related to planning and task instruction (egocentric vs. allocentric reach) on the integration of target and landmark information into the cortical reach network—through the application of graph theoretical analysis (GTA) on fMRI data (Chen et al. 2014). Importantly, there was no difference in the stimuli presented in these tasks, but only the instruction to ignore or use landmarks for coding reach location (Musa et al. 2024). We hypothesized that the *Allocentric* instruction would induce greater global network connectivity, specifically through the integration of dorsal and ventral visual pathways and recruitment of additional Hubs. In addition, we explored the role of global activation trends in interregional cortical communication, by removing these trends from our analysis. Our results suggest that global activation trends related to reach planning reduced global synchrony while enhancing clustering in a reduced number of modules. Further, whereas the egocentric instruction resulted in four functional cortical modules with Hubs clustered in visual and sensorimotor cortex, the allocentric instruction resulted in the conjoining of nodes in ventral visual cortex with the ‘dorsal stream’ module 1, and the recruitment of additional hubs in ventral-posterior and frontal cortex areas associated with allocentric coding and ego-allocentric integration respectively. Finally, our decoding results confirmed that modularity, especially in modules 2 and 3, was the most important parameter for distinguishing the influence of task instruction on network properties. These findings suggest that functional network properties are important for understanding the influence of task instruction on the use of sensory stimuli for visuospatial and motor processing.

### 4.1 Influence of Global Trends on Network ‘Connectivity’

Planning and execution of reaches results not only the recruitment of sensorimotor cortex, but also progressive, widespread cortical activation (Medendorp et al., 2003; Cappadocia et al. 2017), including reactivation of early visual cortex (Gallivan & Culham, 2015; Blohm et al., 2019; Velji-Ibrahim et al., 2022; Monaco et al., 2024). Whereas the specific sensorimotor recruitment and sensory reactivation aspects have received much attention, the general activation trends are often ignored or dismissed as arousal, preparation, or perhaps just an artifact of cumulative BOLD activation. Here we investigated if these trends carry meaningful information, specifically for the integration of local signals across cortex.

When global trends were removed from the BOLD signal, we observed significant changes in network parameters, particularly global synchrony, which is thought to be an important mechanism for ‘binding’ of oscillating signals between different regions (Singer, 1993, 1999; Miller et al., 2012; Blohm et al., 2019). While our fMRI data cannot support or contradict those specific mechanism, they are consistent with the general notion that global trends play a role in maintaining large-scale network synchronization and align with the hypothesis that global BOLD trends reflect long-range communication across brain regions, which can influence local network dynamics (Smith et al., 2012). Removing global trends also increased global clustering, and reduced modularity, reduced number of modules in the Allocentric network and decreased integration of regions traditionally involved in dorsal and ventral stream processing (Figure 4). The latter highlights the importance of baseline brain activity in maintaining network integrity across brain regions (Raichle et al., 2001). Finally, detrending improved task classification based on modularity. These results suggests that removal of such trends is a useful tool to identify task-specific functional segregation.

### 4.2 Global Network Parameters and Behavior

We found that global network features, such as clustering coefficient, global efficiency, and energy, were not particularly discriminative for the task conditions. This suggests that global connectivity measures may be too coarse to capture the subtle, task-specific differences in network organization that underlie spatial goal encoding. In contrast, cross-validated decoding methods demonstrated that task-specific differences could be reliably predicted using features derived from network modularity, even when global parameters showed no significant task-related differences. This is consistent with previous studies, which have suggested that network modularity—an intermediate-level feature that reflects the segregation of brain regions into functionally specialized clusters—is a more sensitive marker for distinguishing between cognitive states (Bassett & Sporns, 2017). These findings emphasize the importance of examining network modularity when investigating the influence of cognitive tasks, as it provides a more nuanced understanding of brain function than broad global parameters alone. In the following sections we examine the influence of task instruction on our network modules and hubs.

### 4.3 The Egocentric Network

#### Egocentric Modules

As noted above, modularity was an important factor in determining the influence of task instruction on our functional networks. The egocentric instruction resulted in four primary modules (at the standard 1.0 resolution threshold), suggesting that the brain uses specialized clusters regions for processing sensory information for action (Figure 4). The superior occipital-parietal / somatomotor module (1) clearly corresponds to the dorsal visual stream and areas involved in sensorimotor transformations (Andersen & Buneo, 2002; Iacoboni, 2006; Gallivan & Culham, 2015; Vesia & Crawford, 2012; Monaco et al., 2024). The inferior parietal / lateral frontal module (2) includes areas with high level spatial processing and top-down instructions (Silver & Kastner, 2009; Peers et al., 2005). The superior temporal / inferior frontal module (3) appears to span auditory cortex and insula so might be involved in multisensory processing of the instruction (Bamiou et al., 2003; Foxe et al., 2002). The inferior occipital-temporal / prefrontal module (4) incorporates elements of the ventral visual stream (Conway, 2018; Kravitz et al, 2011, 2013) and hippocampus, which are involved in configurational computations and spatial memory (Poucet, 1993) and communicate with frontal cortex for object recognition, and the storage and retrieval of spatial memory configurations in the presence of cues (Kar et al., 2019; Kar & DiCarolo, 2021; Zorzo et al., 2021).

#### Egocentric Hubs

Local hubs (Figure 7A) were primarily located in posterior regions, such as the superior parietal lobule and post-central cortex, which are integral for visuomotor processing (Moro et al., 2017). These hubs likely serve as critical nodes for integrating sensory input with motor output. Global hubs (Figure 8A) are areas that possibly play a role in coordinating activity across the different Egocentric network modules described above. These included regions such as the ventral extrastriate cortex, superior extrastriate cortex, and prefrontal cortex—areas known to support visual processing and decision-making in visuomotor tasks (Goodale & Milner, 1992; Pelli et al., 2006).

### 4.4 Changes Observed in The Allocentric Network

Overall, the Allocentric task produced a more integrated network with fewer modules, but more hubs compared to the Egocentric task. A major change observed in the Allocentric task, compared to Egocentric, was a reduction from four to three modules (at the standard 1.0 resolution factor). In part this was due to a conjoining of the ventral occipital (module 4) and dorsal visual pathways (module 1), which are typically thought to support egocentric vs. allocentric functions respectively (Goodale & Milner, 1992). This integration is consistent with the computational need to 1) compute configurational information in the ventral stream, i.e. target relative to landmark (Chen et al. 2014; 2020), and 2) incorporate this information into visuomotor processes typically associated with the dorsal stream.

However, it is equally important that the more anterior temporal portions of egocentric module 4 joined module 2 in the allocentric task, including areas such as auditory cortex and insula. The appears to be consistent with the need for high-level sensory integration in this task, such as relating the instruction to specific visual configurations. Further, our classification analysis (Fig. 10) showed that modules 2 and 3 -conjoining temporal and inferior parietal cortex to prefrontal cortex, were most important for discriminating task. This likely relates to the more linguistic and conceptual aspects of understanding the task, and the role of prefrontal cortex in cognitive control of action (Miller, 2000; Ridderinkhof et al, 2004). Consistent with this interpretation, some regions within these modules (frontal cortex, temporal-parietal junction) have been also implicated in egocentric / allocentric switching in a spatial memory task (Orti et al. 2024).

A direct comparison of ‘hubness’ between tasks (Figure 9) revealed widespread increases in the Allocentric task (Figure 9), especially in Eigenvector Centrality. These changes spanned nodes within previously reported networks including the LimbicB, SalVentAttnA, ContA, DefaultA, DefaultB, SomMotA, TempPar, and VisPeri (Schaefer et al., 2018). These findings align with previous research suggesting that the Allocentric task places higher demands on cognitive control and requires greater interaction between sensory, memory, and motor networks (McNamara & Shelton, 2000).

The specific Eigenvector hubs for the Allocentric task (Fig. 7B) correspond to areas in the control, salience and dorsal attention networks (Schaefer et al., 2018). The presence of additional Eigenvector hubs in extrastriate cortex, especially the inferior areas, seems consistent with the increased visual complexity of this task. The presence of additional hubs in prefrontal cortex could reflect the need for increased top-down cognitive demands of the Allocentric task, which likely requires greater executive control, attentional allocation, and memory integration (Desimone & Duncan, 1995). Further, some of these regions (e.g., frontal eye fields) have been shown to play a role in egocentric-allocentric integration (Chen et al., 2018; Bharmauria et al., 2020, 2021; Schutz et al., 2023). Overall, these areas seem likely to support this particular task. Likewise, the higher number of betweenness centrality hubs in the Allocentric task (Figure 8B) compared to egocentric (8A) supports the greater need for communication between modules in this task.

### 4.5 Lateralization

While examining hemispheric asymmetry in the networks, we found that both the Egocentric and Allocentric networks exhibited substantial bilateral symmetry in terms of module organization and hub locations. Allocentric ‘Hubs’ which showed increased centrality in terms of Eigenvector and Betweenness Centrality (Figure 9) were areas that are integral to visual processing and spatial awareness, particularly in the context of landmark-based navigation (Bremmer et al., 2001; Grefkes & Fink, 2005). Lateralization in the Allocentric network was observed, with key regions showing hemispheric dominance. The left hemisphere was associated with regions such as the orbitofrontal cortex (OFC), parietal operculum (ParOper), and dorsolateral prefrontal cortex (PFCd), while the right hemisphere showed significant involvement of the insula (Ins), inferior parietal lobule (IPL), and somatomotor areas (SomMotA). The left hemisphere regions are primarily associated with decision-making and motor planning, while right hemisphere regions are involved in sensory integration and spatial processing.

### 4.6 Possible Physiological Basis of Network

The BOLD signals observed in this study reflect large-scale population-level neural activity, likely driven by synaptic inputs from various brain regions (Logothetis et al., 2001). Numerous physiological studies have observed configurational information (feature vs. feature, object vs. object) in the ventral visual stream, including areas possibly corresponding to ExStr_1 and ExStrInf_2 in our study (Kravitz et al., 2013s). Fewer studies have examined target-landmark interactions for action, it has been shown that target and landmark signals coexist and interact to produce landmark-centred codes in frontal and supplementary cortex (Schutz et al., 2023). Further Bharmauria et al. (2020, 2021) showed that landmark shifts in these areas are detected in the memory responses and integrate in the motor responses of these structures. Finally, Taghizadeh et al. (2024) found evidence for both egocentric and allocentric coding in dorsal stream reach areas of monkey cortex, corresponding to our areas PC_3, SomMotA_1 and SomMotA_4. These studies suggest that our networks reflect real physiological processes and conversely our results suggest that simultaneous physiological recordings over a broad range of areas are likely necessary to fully understand how these signals are integrated at a cellular level.

### 4.7 General Implications

The findings from this study suggest that the modular organization of brain networks, particularly in terms of how regions segregate and integrate information, plays a critical role in supporting complex cognitive behaviors such as spatial goal encoding. Specifically, we find that task-related instructions can have a profound influence on modularity and hubness, likely influencing the way the brain processes sensory information (which was otherwise equalized in our two tasks). Further, it appears that widespread activation attributed to recurrent motor signals (Blohm et al. 2019) may play a complementary role in integrating these signals across cortex when it is time for action. These results could have broad implications for understanding how brain signals are organized itself to handle other types of complex, goal-directed behaviors. Furthermore, the ability to decode task-specific brain networks based on modularity measures suggests that event-related connectivity analysis could serve as a powerful tool for clinical diagnostics. For example, disruptions in the modular structure of brain networks may be indicative of neurological disorders that affect cognitive function, such as Alzheimer’s disease or schizophrenia (Bassett et al., 2011) or explain why visual landmarks benefit movement initiation for Parkinsons’s patients (Russo et al., 2022).

### 4.8 Conclusions

In conclusion, we have identified complementary mechanisms for segregating and integrating networks in response to task instructions: 1) topological segregation of similar signals into functional modules for visually-evoked movements, 2) integration across modules for specific task (such as the dorsoventral and posterior-anterior integration of modules observed in our allocentric task), 3) recruitment of specialized hubs for specific task requirements, and 4) promotion of global synchrony through the overall activation trends that emerge during movement planning. Although these results emerge from a very specific dataset, we suggest that they would likely generalize to other complex visually-guided behaviors, and that they would degrade during disease processes.

## Ethics Approval

Investigating cortical mechansims underlying visuomotor control using functional Magnetic Imaging (fMRI) in humans [EthicsProtocol #3730]

Certificate #: e2023-336

## Author Contributions

LM: Conceptualization, Data Curation, Analysis, Visualization, Writing – Original Draft

AG: Supervision (Graph Theory Analysis)

YC: Investigation (Data Collection), Data Curation

JDC: Conceptualization, Funding Acquisition, Project Administration, Supervision, Writing – Review and Editing

## 5. Funding

This work was funded by the Canadian Institutes for Health Research [grant number MOP-68812] and the Vision: Science to Applications (VISTA) Program [grant number 102001171]. L. Musa and A. H. Ghaderi were supported by VISTA. J.D. Crawford was supported by a Canada Research Chair, Canada First Research Excellence fund [grant number 101035774].

## Supplementary Figures

**Supplementary Figure 1.**
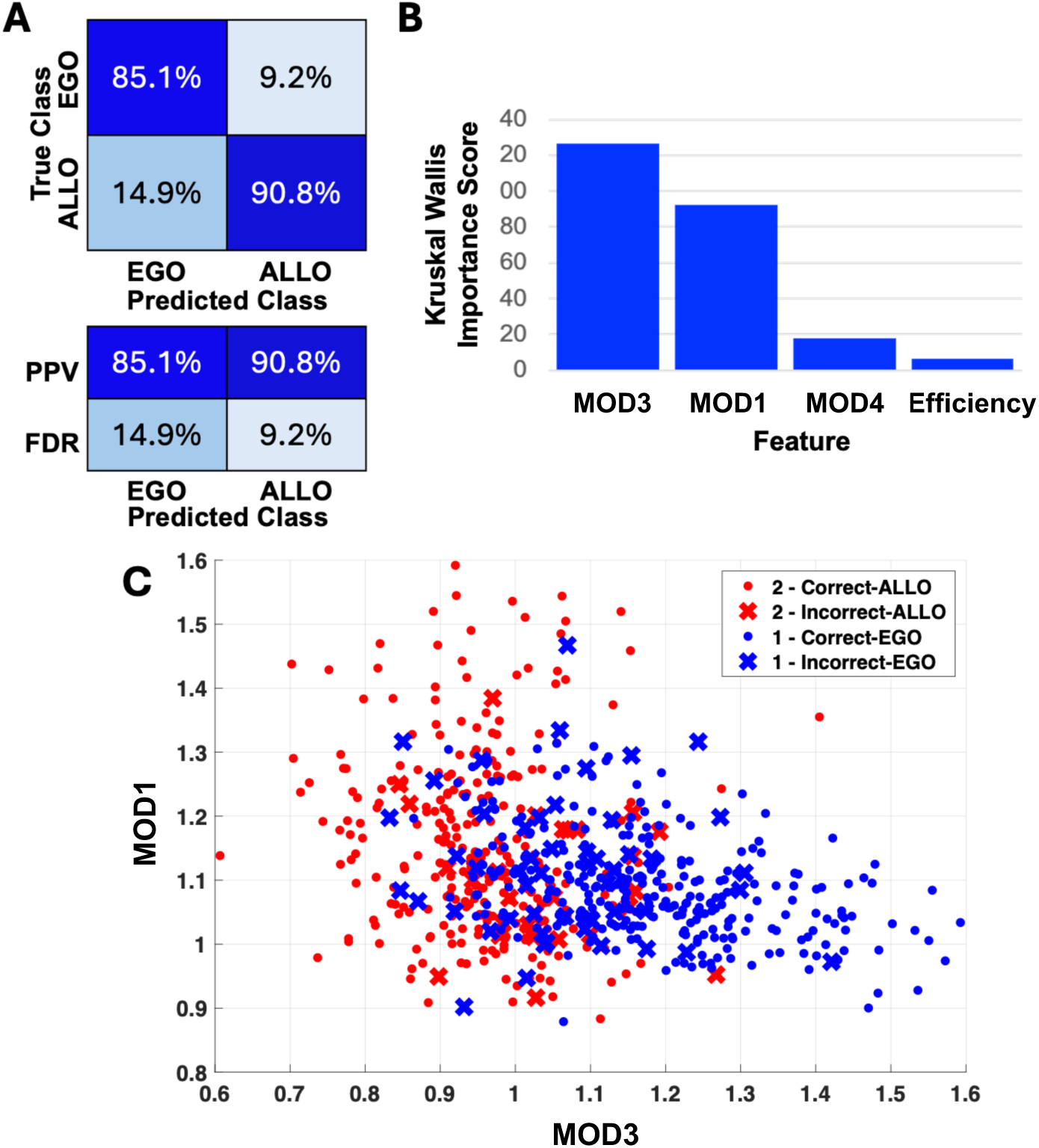
Classification Analysis. Performance of the SVM classifier in distinguishing between the Egocentric and Allocentric networks in the detrended network (A) The confusion matrix (top panel table) comparing the percentage of data points in the four categories shown, top row: data points with a true and predicted class of Egocentric (EGO (top left cell), data points with a true class of Egocentric (EGO) and predicted class of Allocentric (ALLO) (top left cell), and the corresponding categories for the Allocentric task in the second row. The bottom panel table shows the PPV and FDR of the Egocentric network (EGO, left column) and the Allocentric network (ALLO, right column). (B) Importance scores of the features classifying Egocentric and Allocentric networks in the original dataset, as determined by the Kruskal-Wallis approach. (C) Scatter plot of distribution of datapoints predicted across two dimensions: (modularity score of module 3 [MOD3] vs modularity score of module 1 [MOD1]) out of the total four classifier dimensions. Correct classifications are shown as dots whereas misclassifications are shown as crosses. The two networks being classified, Egocentric and Allocentric are shown in blue and red, respectively.

## Notes

### Competing Interest Statement

The authors have declared no competing interest.

